# Spatiotemporal atlas resolves the mesenchymal-epithelial stem cell axis governing human hair follicle aging and pathology

**DOI:** 10.64898/2026.09.22.753364

**Authors:** Xiaoyu Wei, Yanwen Xu, Jiaxin Du, Ruikang Li, Zhentao Zhou, Yeya Yu, Zishuo Yuan, Hanxiao Cheng, Yujia Jiang, Fei Zhu, Pengfei Cai, Haiyan Shen, Shuai Wang, Yue Yuan, Xiawei Liu, Tao Yang, Yinghua Huang, Yiwei Lai, Shiwei Wang, Hanbo Li, Jun Xie, Ying Gu, Dong Niu, Zhenxing Wang, Xun Xu, Longqi Liu, Mingxing Lei, Pengcheng Guo, Jufang Zhang

**Affiliations:** State Key Laboratory of Genome and Multi-omics Technologies, BGI Research, Hangzhou 310030, China; Key Laboratory of Brain Cell Mapping of Zhejiang Province, BGI Research, Hangzhou 310030, China; Department of Medical Cosmetic Center, Affiliated Hangzhou First People’s Hospital, School of Medicine, Westlake University, Hangzhou 310006, China; BGI Research, Hangzhou 310030, China; College of Life Sciences, University of Chinese Academy of Sciences, Beijing 100049, China; Key Laboratory of Resources Biology and Biotechnology in Western China, Ministry of Education, Provincial Key Laboratory of Biotechnology of Shaanxi Province, the College of Life Sciences, Northwest University, Xi’an 710069, China; Key Laboratory of Spatial Omics of Zhejiang Province, BGI Research, Hangzhou 310030, China; BGI Research, Shenzhen 518083, China; Department of Biology, University of Copenhagen, DK-2100 Copenhagen, Denmark; BGI Research, Qingdao 266555, China; China National GeneBank, BGI Research, Shenzhen 518120, China; Guangdong Provincial Genomics Data Center, BGl Research, Shenzhen 518120, China; BGI Cell, Hangzhou 310030, China; MOE Key Laboratory of Coal Environmental Pathogenicity and Prevention, Shanxi Medical University, Taiyuan 030001, China; Shanxi Medical University-BGI Collaborative Center for Future Medicine, Shanxi Medical University, Taiyuan 030001, China; BGI Cell, Shenzhen 518083, China; College of Animal Science and Technology, College of Veterinary Medicine, Zhejiang A&F University, Hangzhou 311300, China; Department of Plastic Surgery, Union Hospital, Tongji Medical College, Huazhong University of Science and Technology, Wuhan, 430022, China; Key Laboratory of Biorheological Science and Technology of Ministry of Education and 111 Project Laboratory of Biomechanics and Tissue Repair, College of Bioengineering, Chongqing University, Chongqing 400044, China; Center for Plastic & Reconstructive Surgery, Department of Plastic & Reconstructive Surgery, Zhejiang Provincial People’s Hospital (Affiliated People’s Hospital), Hangzhou Medical College, Hangzhou 310006, Zhejiang, China

## Abstract

How adult stem cell populations coordinate tissue regeneration across physiological and pathological states remains unclear. Hair follicles contain epithelial and mesenchymal stem cells that codetermine hair regeneration. Here, we integrate a multimodal spatiotemporal atlas with functional validation to define the mesenchymal-epithelial stem cell dynamics governing human hair follicle homeostasis, aging, and androgenetic alopecia. Aging is characterized by passive depletion of hDSC-derived regenerative cues, whereas AGA involves a shift of hDSC toward an inhibitory signaling hub that promotes HFSC dysfunction. We further pinpoint hDSC-secreted PTN as a key regenerative factor that promotes hair growth and counteracts androgen-induced growth suppression. This study provides a holistic framework of adult stem cell crosstalk across physiological and pathological states, offering a mechanistic roadmap to reinstate tissue regeneration.

## Introduction

Adult stem cells are the fundamental units of tissue homeostasis and organ regeneration, possessing the unique capacity to self-renew and generate specialized progeny across the human lifespan^1-4^. The hair follicle, a complex mini-organ that undergoes lifelong cyclical renewal, serves as an ideal paradigm for decoding the regulatory logic of human stem cell biology^5-8^. This exceptional regenerative potential is driven by two distinct stem cell populations residing in specialized niches: epithelial hair follicle stem cells (HFSC), which reside in the bulge and generate the diverse epithelial lineages of the hair shaft and surrounding sheath^9-11^, and mesenchymal hair follicle dermal stem cells (hfDSC), which occupy the dermal cup and replenish the dermal papilla (DP) and dermal sheath (DS)^12-14^. Although the lineage-specific differentiation programs of these populations are well-established and mesenchymal-epithelial crosstalk is universally recognized as a fundamental driver of hair follicle renewal, it remains largely elusive whether these distinct mesenchymal and epithelial stem cell pools engage in direct spatiotemporal interactions to coordinately regulate the regenerative process^9,11,12,15^.

The established paradigm of hair follicle biology posits that the DP serves as the primary inductive center required to trigger telogen-to-anagen transition by instructing HFSC to exit quiescence^6,15^. However, this signaling model must contend with a significant spatial challenge inherent to the hair cycle^7,11,15,16^. As the follicle undergoes regenerative growth, the resulting anatomical remodeling causes a dramatic segregation of the niche, transporting the DP deep into the dermis and far away from the bulge-resident HFSC^5,17,18^. This physical distance is particularly striking in human follicles, where the growth phase can persist for several years, raising fundamental questions regarding how the stable and continuous signals necessary for sustained HFSC activation are reliably maintained across this expanded tissue architecture to ensure long-term niche homeostasis and uninterrupted hair production^19^. We propose that the functional maintenance of the follicle relies on a direct mesenchymal-epithelial stem cell axis, where the parent mesenchymal stem cell pool (hfDSC) serves as a critical signaling hub to bridge this spatial gap and synchronize niche dynamics throughout the hair cycle.

The homeostatic balance of this multi-lineage niche is significantly disrupted during physiological aging and in prevalent hair loss disorders such as Androgenetic Alopecia (AGA)^11,20-22^. These conditions drive a hallmark transition toward progressive follicular miniaturization and a pathological shift into a prolonged telogen state. While focused investigations into individual stem cell lineages have provided foundational insights into the mechanisms of hair follicle regeneration, aging and AGA, the complex synergistic orchestration between distinct stem cell pools remains poorly explored^13,23-25^. Furthermore, although recent single-cell transcriptomic studies have provided valuable snapshots of cellular heterogeneity in the human hair follicle, the precise identity and human-specific markers of hair follicle dermal stem cells (hDSC) have remained elusive^23,24,26-29^. Consequently, these approaches often lack the spatial coordinates and molecular resolution required to decipher the interfacial communication between the parent mesenchymal and epithelial stem cell pools.

To address these gaps, we generated a comprehensive multimodal atlas of the human scalp, integrating high-resolution Stereo-seq^30^, scRNA-seq, and scATAC-seq data from a well-characterized cohort spanning the human lifespan and including individuals with AGA. By resolving the molecular architecture of the hair follicle at single-cell and spatial resolution, we identify hDSC as a major spatially adjacent signaling hub. We reveal that hair follicle aging and AGA are associated with disruption of this mesenchymal-epithelial stem cell axis, characterized by passive depletion in aging and active pathogenic transformation in AGA. Elucidating the dynamics of this axis reveals a higher-order regulatory logic of tissue regeneration, suggesting that the functional integrity of complex organs is predicated on the reciprocal synergy between distinct stem cell populations rather than isolated lineage behaviors. Finally, we identified and functionally validated shared molecular drivers across both physiological and pathological decline, providing candidate strategies for restoring regenerative signaling.

## Results

### Multimodal spatiotemporal atlas of human scalp

To dissect the spatiotemporal dynamics within human hair follicles during aging and AGA, we conducted multimodal single-cell and spatial transcriptomics analyses on human scalp tissues. We obtained samples from: 1) discarded occipital scalp tissues from 22 healthy male volunteers (HO) aged 3 to 77 years, stratified into four age groups (Juvenile: 3 to 17 years, prime age: 18 to 39 years, middle age: 40 to 59 years, and old age: 60 to 77 years), and 2) occipital (hair-bearing, AO) and frontal (hairless, AF) scalp tissues from 6 male AGA patients diagnosed as levels M2 and M3 based on the BASP classification system **(Fig. 1a and Extended Data Fig. 1a-c)**. The scRNA-seq dataset comprised 262,809 cells from 30 samples (22 HO and 4 AO, 4 AF from 4 AGA patients), with a median of 4,765 unique molecular identifiers (UMIs) and 1,838 genes per cell **(Extended Data Fig. 1d and Supplementary Table 1)**. The scATAC-seq dataset included 91,223 cells from 12 samples (9 HO and 4 AO, 3 AF from 4 AGA patients), with a median of 11,489 fragments and a TSS enrichment score of 5.05 per cell **(Extended Data Fig. 1e and Supplementary Table 2)**. For the Stereo-seq, we applied cell segmentation based on nucleic acid staining images^31^ for all slides from 23 samples (16 HO and 4 AO, 3 AF from 4 AGA patients) to generate image-guided gene by cell matrix containing 2,124,467 segmented cells **(Extended Data Fig. 1f and Supplementary Table 3)**.

**Fig. 1.**
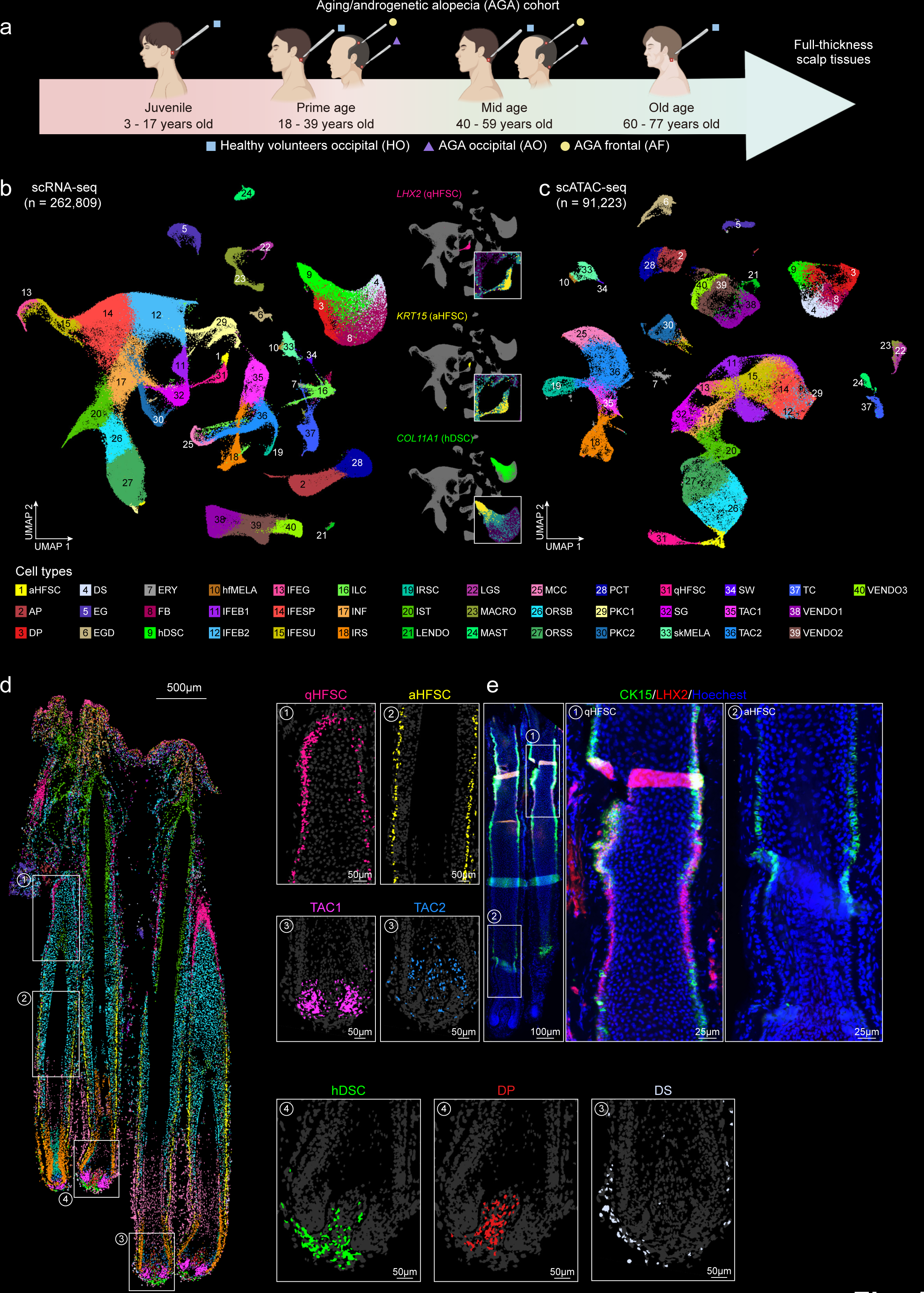
Spatiotemporal multi-omics atlas of the human scalp in aging and AGA. **a,** Schematic of the experimental design. Full-thickness human scalp samples were obtained from male healthy donors and patients with AGA. Samples were processed for scRNA-seq, scATAC-seq, and Stereo-seq. Healthy donor samples, obtained from the occipital scalp (HO), were stratified into four age groups: Juvenile (3 - 17 years old), prime age (18 - 39 years old), mid age (40 - 59 years old), and old age (60 - 77 years old). AGA patient samples were obtained from both occipital and frontal scalps. HO, healthy occipital; AO, AGA occipital; AF, AGA frontal. **b,** Uniform Manifold Approximation and Projection (UMAP) of single-cell profiles. Left: UMAP visualization of 40 distinct cell types identified via scRNA-seq (n = 262,809). Right: Targeted UMAP highlighting 3 adult stem cell populations (qHFSC, aHFSC, and hDSC) with feature plots showing the expression of canonical markers (*LHX2*, *KRT15*, and *COL11A1*). aHFSC, actived hair follicle stem cell; AP, arrector pili cell; DP, dermal papilla cell; DS, dermal sheath cell; EG, eccrine gland cell; EGD, eccrine gland duct cell; ERY, erythrocyte; FB, fibroblast; INF, infundibular cell; IRS, inner root sheath cell; hDSC, human hair follicle dermal stem cell; hfMELA, hair follicle melanocyte; IFEB1, interfollicular epidermis basal cell 1; IFEB2, interfollicular epidermis basal cell 2; IFEG, interfollicular epidermis granular cell; IFESP, interfollicular epidermis spinous cell; IFESU, interfollicular epidermis suprabasal cell; ILC, innate lymphoid cell; IRSC, inner root sheath cuticle cell; IST, isthmus cell; LGS, Langerhans cell; LENDO, lymphatic endothelial cell; MACRO, macrophage; MAST, mast cell; MCC, medulla cortex cuticle cell; ORSB, outer root sheath basal cell; ORSS, outer root sheath suprabasal cell; PCT, pericyte; PKC1, proliferating keratinocyte 1; PKC2, proliferating keratinocyte 2; qHFSC, quiescent hair follicle stem cell; SG, sebaceous gland cell; TC, T cell; TAC1, transit amplifying cell 1; TAC2, transit amplifying cell 2; skMELA, skin melanocyte; SW, Schwann cell; VENDO1, vascular endothelium cell 1; VENDO2, vascular endothelium cell 2; VENDO3, vascular endothelium cell 3. **c,** UMAP of 39 cell types defined by scATAC-seq (n = 91,223). **d,** Spatial distribution and molecular validation of cell types in the hair follicle. Left: Spatial visualization of 40 cell types in a representative Stereo-seq section, deconvoluted by RCTD. White boxes indicate regions magnified in right panels. Right: Magnified views illustrating the spatial coordination of selected cell types. **e,** Immunofluorescence co-staining of CK15 (green) and LHX2 (red) in human hair follicles. Right: Highlight of the distinct anatomical niches of qHFSCs (left) and aHFSCs (right). Scale bar, 50 μm.

Integration of scRNA-seq data from all samples revealed a high degree of correlation across age groups, AGA status, and scalp regions, indicating minimal cell cluster bias **(Fig. 1b and Extended Data Fig. 2a)**. Uniform manifold approximation and projection (UMAP) visualization of the scRNA-seq data enabled robust identification of 40 distinct cell types or subtypes within human scalp based on the known marker genes **(Fig. 1b, Extended Data Fig. 2b and Supplementary Table 4)**. We identified several interfollicular epithelial cell types residing in distinct skin layers, including interfollicular epidermis basal cells (IFEB1 and IFEB2), interfollicular epidermis spinous cells (IFESP), interfollicular epidermis suprabasal cells (IFESU), interfollicular epidermis granular cells (IFEG), and skin melanocytes (skMELA). Within the hair follicle, the major epithelial cell types were medulla cortex cuticle cells (MCC), inner root sheath cells (IRS), inner root sheath cuticle cells (IRSC), outer root sheath basal cells (ORSB), outer root sheath suprabasal cells (ORSS), transit amplifying cells (TAC1 and TAC2), isthmus cells (IST), infundibular cells (INF), hair follicle melanocyte (hfMELA), and hair follicle stem cells (HFSC). Notably, we were able to distinguish between activated HFSC (aHFSC), characterized by high expression of *LGR5, RUNX1,* and *ANGPTL7*^10,32^, and quiescent HFSC (qHFSC), characterized by high expression of *NFATC1, CXCL14* and *KRT15*^33,34^. Within the hair follicle’s mesenchymal compartment, alongside canonical dermal papilla (DP) and dermal sheath (DS) cells, we identified a distinct subcluster characterized by high expression of human hair follicle dermal cup markers *RBP4, MMP11,* and *ASPN*^35^. In addition, our dataset captured the dermal fibroblasts (FB), marked by *APOD*, *CXCL12*, and *GSN*^28^, as well as diverse immune cell types, including macrophages (MACRO), Langerhans cells (LGS), innate lymphoid cells (ILC), T cells (TC), and mast cells (MAST). We also identified other cell types within the hair follicle niche, including 3 types of vascular endothelial cells (VENDO1, VENDO2, VENDO3) and lymphatic endothelial cells (LENDO), Schwann cells (SW), and gland-related cells, such as eccrine gland cells (EG), eccrine gland duct cells (EGD), sebaceous gland cells (SG), and arrector pili cells (AP), erythrocytes (ERY), fibroblast (FB), pericytes (PCT), and proliferating keratinocyte cells (PKC1, PKC2). Analysis of the scATAC-seq data also showed robust identification of the main cell types with a high degree of correlation across sample conditions **(Fig. 1c and Extended Data Fig. 2c)**.

To visualize the spatial organization of diverse cell types within different anatomic regions at the whole hair follicle scale, we projected scRNA-seq data onto the Stereo-seq gene-by-cell matrix using RCTD^36^ to determine the spatial distribution of each cell type. Our spatial map revealed distinct localization patterns for all 40 identified cell types across all Stereo-seq sections, and hair growth phases (anagen, telogen) and miniaturization were determined by hematoxylin-eosin (H&E) staining of adjacent sections **(Fig. 1d, Extended Data Fig. 2d and Extended Data Fig. 3-5)**^37^. This multimodal map confirmed the expected zonation of the epithelial lineages, with qHFSC (CK15^+^LHX2^+^) were predominantly located in the bulge region, aHFSC (CK15+LHX2^-^) positioned immediately below them, and their progeny TAC1 and TAC2 progressively shifting downward along the follicular axis **(Fig. 1d-e)**. Critically, this spatial analysis provided the definitive evidence required to resolve the identity of the ambiguous mesenchymal subcluster. DP and DS cells mapped to their canonical anatomical locations, and the subcluster expressing dermal cup markers was specifically and exclusively restricted to the dermal cup, which represents the established anatomical niche for murine hair follicle dermal stem cells (hfDSC) **(Fig. 1d)**^12^. Based on this definitive spatial evidence and its unique transcriptional profile, we designated this population as human Hair Follicle Dermal Stem Cells (hDSC). Although hDSC occupy the same anatomical niche as murine hfDSC, our comparative analyses revealed that they exhibit profound species-specific transcriptional divergence **(Extended Data Fig. 5a-d)**. This integrated, multimodal approach thus enabled the definitive identification and spatial localization of a putative stem cell population within the adult human hair follicle’s dermal niche.

We have generated a comprehensive single cell multi-omics spatiotemporal atlas of human scalp, comprising over 2.4 million cells, that encompasses both physiological aging and AGA progression. We spatially resolved the main cell types and subtypes within the hair follicle, including the identification of hDSC, aHFSC and qHFSC, providing a valuable resource for studying hair follicle biology. The data are accessible via the interactive data portal at https://db.cngb.org/stomics/hhaamstar/.

### Putative stem cell lineage transitions during hair follicle regeneration

A defining characteristic of somatic stem cells is their ability to reconstitute all cell types within their tissue of origin. Hair follicles harbor diverse somatic stem cell populations, with both HFSC and hDSC playing crucial roles in the cyclical bouts of hair regeneration. Lineage tracing and scRNA-seq studies in rodent models have illuminated the distinct differentiation trajectories of epithelial and mesenchymal cells within the hair follicle^12,33^. However, these processes remain poorly characterized in humans. To address this, we employed the Dynamo^38^ algorithm for the Stereo-seq data and Monocle 3^39^ for the scRNA-seq data to predict cell lineage transitions within these two compartments. Our findings corroborate the proposed lineage trajectory in which qHFSC activate and migrate downward to become aHFSC, which subsequently progress through TAC1 and ORSB to generate the IRS and ORSS, respectively^11,33^. Additionally, our model suggested that TAC1 cells can transiently differentiate into TAC2 that give rise to both MCC and IRSC **(Fig. 2a-b and Extended Data Fig. 6a)**^40^. Parallel analysis of the mesenchymal niche delineated a conserved trajectory in which hDSC serve as the source for both DP and DS compartments **(Fig. 2c and Extended Data Fig. 6b-c)**^12,13^. However, a cross-species comparison highlights that despite the conserved directional flow, the transcriptional programs governing human hDSC-to-DP/DS differentiation are markedly distinct from those in mice, underscoring the necessity of human-specific molecular maps **(Extended Data Fig. 6d-e)**.

**Fig. 2.**
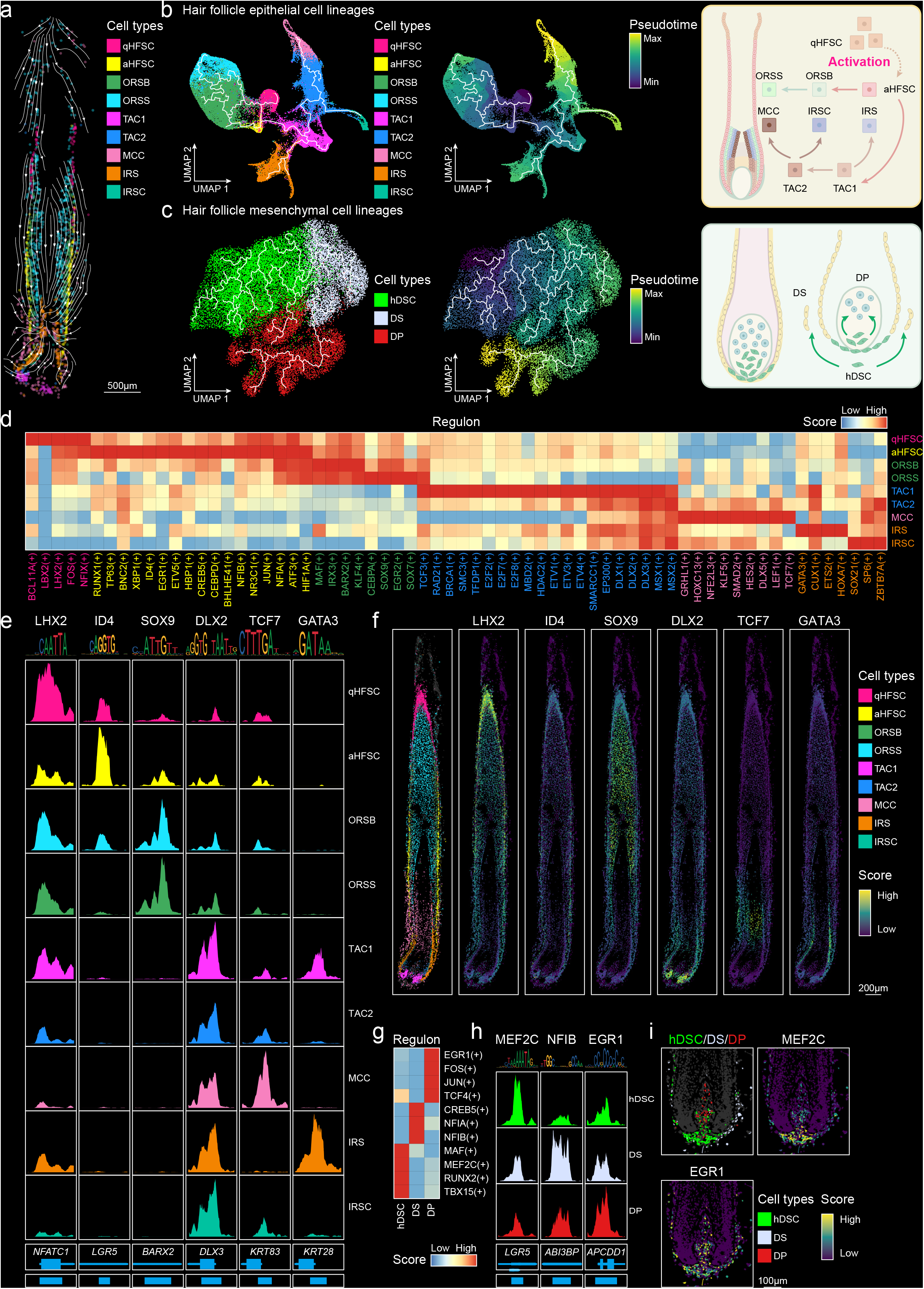
Putative cell lineage transitions during hair follicle regeneration. **a,** RNA velocity streamline plot showing the predicted differentiation trajectories of hair follicle epithelial cells in a representative Stereo-seq section. **b,** Pseudotime analysis of hair follicle epithelial cell lineages in the scRNA-seq dataset using Monocle 3. Dots are colored by cell types (left) or pseudotime (middle). Schematic representation of the stepwise differentiation trajectories of hair follicle epithelial cells (right). **c,** Pseudotime analysis of hair follicle mesenchymal cell lineages in scRNA-seq dataset using Monocle 3. Dots are colored by cell types (left) or pseudotime (middle). Schematic representation of the differentiation trajectories of hair follicle mesenchymal cells (right). **d,** Heatmap showing the average regulon activity in hair follicle epithelial lineages. Color intensity represents the normalized AUC score of the indicated transcription factors (TFs). **e,** Genomic tracks showing chromatin accessibility at representative target gene loci for epithelial lineage specific regulons. Shown from left to right are the *LHX2* locus (qHFSC-specific ATF3 regulon), the *LGR5* locus (aHFSC-specific ID4 regulon), the *BARX2* locus (ORSB/ORSS-specific SOX9 regulon), the *DLX3* locus (TAC1/TAC2-specific DLX2 regulon), the *KRT83* locus (MCC-specific TCF7 regulon), and the *KRT28* locus (IRS/IRSC-specific GATA3 regulon). **f,** Spatial visualization of module scores for target genes corresponding to the epithelial lineage-specific regulons identified in **(e)**. **g,** Heatmap showing the average regulon activity in hair follicle mesenchymal lineages. **h,** Genomic tracks showing chromatin accessibility at representative target gene loci for the hair follicle mesenchymal lineage specific regulons. Left: the *LGR5* locus (hDSC-specific MEF2C regulon); Middle: the *ABI3BP* locus (DS-specific NFIB regulon); Right: the *APCDD1* locus (DP-specific EGR1 regulon). **i,** Spatial visualization of module scores for target genes corresponding to the hair follicle mesenchymal lineage-specific regulons identified in **(h)**.

To dissect the gene regulatory networks governing hair follicle cell identity, we applied SCENIC+^41^ to the epithelial lineage **(Fig. 2d-f, Extended Data Fig. 6f-g and Supplementary Table 5)**. This analysis successfully reconstructed the transcriptional logic governing stemness, activation, and lineage commitment. We first resolved the distinct regulatory networks driving the functional states of the stem cell pool. The quiescent state of qHFSC was supported by the expression of canonical regulators such as *FOXC1* and *NFATC1* **(Extended Data Fig. 6f)**^34^, alongside specifically enriched regulons including BCL11A, LBX2, LHX2, FOS and NFIX, which function to enforce stem cell quiescence and prevent premature activation^42-44^. The activation program was characterized by the robust enrichment of canonical activation driver (RUNX1)^32^ and essential telogen-to-anagen transition regulators TP63 and BNC2^45,46^, operating within a broader regulatory network that includes XBP1, ID4, EGR1, ETV5, HBP1. Crucially, this transition was accompanied by a coordinated metabolic reprogramming module (HIF1A, CREB5, CEBPD, BHLHE41) that orchestrates the bioenergetic shift to promote anaerobic glycolysis required to support the HFSC activation^47^. Collectively, these regulons delineate a highly orchestrated regulatory hierarchy, wherein core identity guardians, quiescence enforcers, and activators act in concert to strictly govern hair follicle stem cell homeostasis and the precision of the hair cycle.

As cells exited the stem cell niche, our analysis resolved their bifurcation into distinct differentiation trajectories. The outer root sheath lineages (ORSB and ORSS) were characterized by a specific suite of regulators, including MAF, IRX3, KLF4, CEBPA, SOX9, EGR2, SOX7, and the canonical marker BARX2^9^. For the inner lineages, cells first entered a highly proliferative Matrix/TAC state. This compartment was characterized by the Wnt signaling effector TCF3^28^ and a coordinated proliferation and priming network encompassing cell cycle drivers (RAD21, BRCA1, SMC3, TFDP1, E2F family), and differentiation-priming factors (ETV1/3/4, YBX1). Furthermore, we identified a core pan-lineage module of transcription factors that exhibit peak expression levels in TACs and maintain broad enrichment across TAC-derived progeny. This module comprises key chromatin modifiers, including MBD2, HDAC2, SMARCC1, and EP300, alongside essential hair follicle differentiation regulators such as DLX1/2/3, MSX1/2, and HES1^9,18,48^. This module likely functions to maintain high levels of chromatin plasticity, providing the necessary epigenetic flexibility for heterogeneous TACs to transition effectively into their respective specialized terminal fates^5^. From this primed pool, terminal differentiation was governed by distinct networks. The Medulla/Cortex/Cuticle (MCC) lineage was defined by key hair shaft regulators (GRHL1, HOXC13, NFE2L3)^9^, integrated with Wnt/β-catenin effectors (LEF1, TCF7)^18^, and KLF5, SMAD2, HES2, DLX5. Concurrently, the Inner Root Sheath (IRS) was governed by established drivers (GATA3, CUX1)^9^, alongside a suite of novel human regulons including ETS2, HOXA7, SOX21, SP6, and ZBTB7A. This comprehensive mapping of the regulatory hierarchy provides a high-resolution framework for understanding the molecular drivers of human hair follicle differentiation.

Applying this validated framework to the mesenchymal lineage, we uncovered the distinct regulatory architectures defining the stem cell and differentiated states **(Fig. 2g-i, Extended Data Fig. 6h-i and Supplementary Table 6)**. We found that the hDSC population is uniquely defined by the activity of regulons including TBX15, RUNX2, MEF2C, and MAF. The terminally differentiated progeny are governed by discrete regulatory programs. The DS population is driven by NFIA, NFIB, and CREB5, whereas the DP is controlled by established factors such as TCF4^49^, alongside JUN, FOS, and EGR1. This analysis demonstrates that the differentiation from hDSC to DP and DS is underpinned by a definitive switch in master-regulatory transcription factor activity, providing mechanistic support for the inferred lineage trajectory.

### Spatially resolved hDSC dysfunction with aging

Our integrated dataset of healthy human scalps spanning four age groups provides a powerful resource to dissect the cellular and molecular mechanisms underlying human hair follicle aging. We first constructed a spatiotemporal landscape of the hair follicle across different ages, revealing pronounced follicle miniaturization with increasing age **(Fig. 3a)**. To determine the susceptible cell types in aging, we correlated cell type proportions with age and identified 9 cell types that decreased and 1 that increased with age (*P* < 0.05, |correlation coefficient| > 0.4) **(Fig. 3b and Extended Data Fig. 8a)**. Notably, hDSC exhibited a marked age-associated reduction, whereas both qHFSC and aHFSC remained relatively stable, consistent with a prior murine study showing that HFSC depletion is a hallmark of pathological rather than physiological aging^50^.

**Fig. 3.**
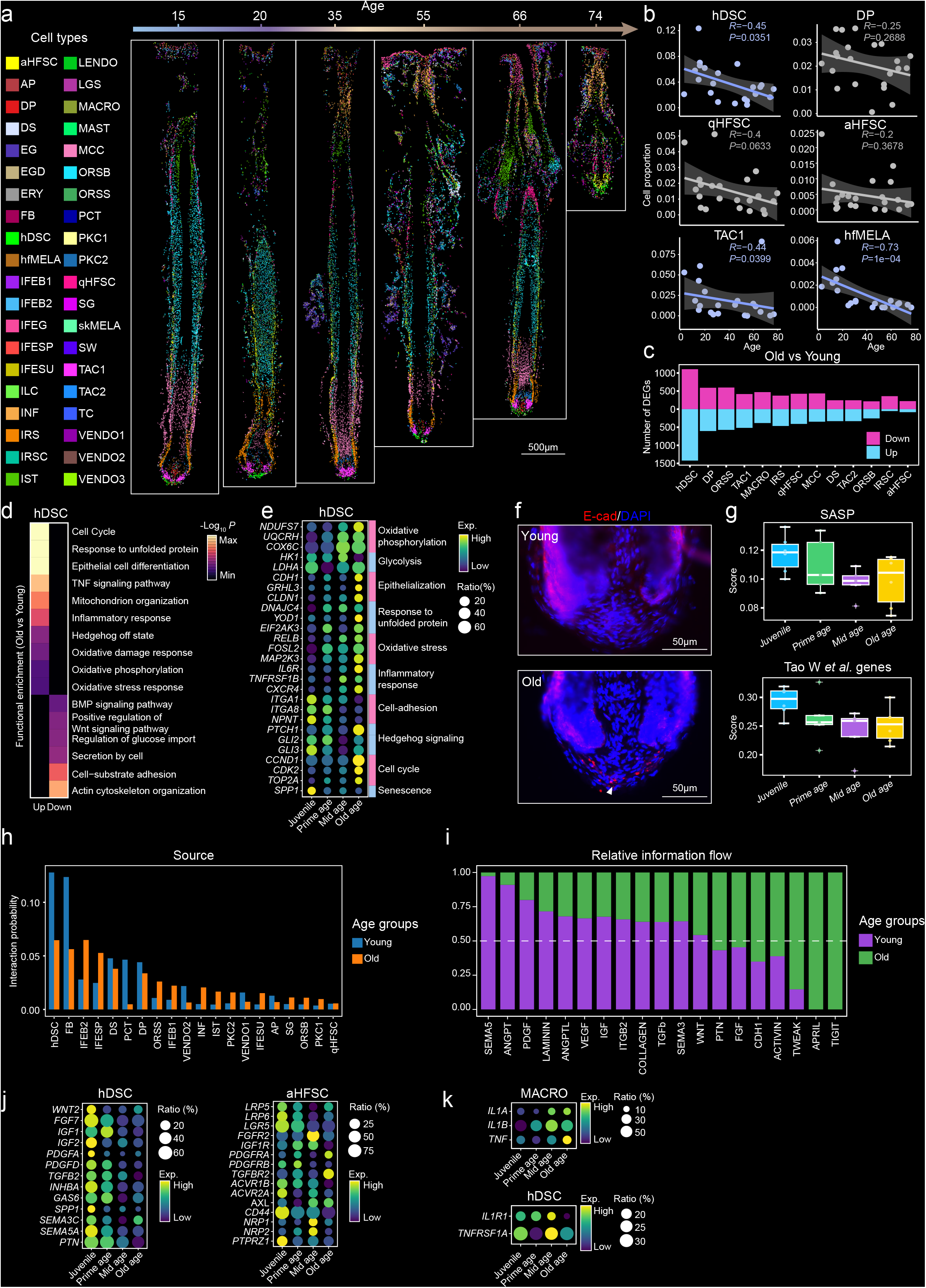
Single-cell Stereo-seq dissects the cellular and molecular dynamics in human hair follicles during aging. **a,** Spatial visualization of cell type distributions in 6 representative Stereo-seq sections from indicated age groups deconvoluted by RCTD and colored by the 40 identified cell types. **b,** Correlation plots showing the relationship between the proportion of indicated cell types and age. *R*, Pearson correlation coefficient; *P*, two-sided P-value; the center represents the average value; shading indicates the 95% confidence interval. **c,** Bar plot showing the number of upregulated (purple) and downregulated (blue) DEGs in indicated cell types in the old group (60-77 years) compared to the young group (12-30 years). **d,** Heatmap of functional enrichment analysis of upregulated and downregulated DEGs in hDSC. Color intensity represents pathway significance (-Log_10_ *P*). **e,** Bubble plot showing the average expression of selected hDSC DEGs related to indicated biological processes in different age groups. Color intensity indicates average gene expression, and dot size represents the percentage of positive cells. **f,** Representative images of immunofluorescence staining for E-cadherin (E-cad) in human scalp sections from young (top) and old (bottom) individuals. Scale bar, 50 μm. **g,** Box plots showing the module scores of senescence-associated secretory phenotype (SASP) genes (left) and aging-related genes taken from Tao *et al*., 2024 (right) in hDSC from indicated age groups. **h,** Bar plot showing the strength of source (ligand) signaling interactions from each cell type in young and old groups, as measured by CellChat. **i,** The sum of differences in interaction probability (relative information flow) for indicated signaling pathways between young and old groups. **j,** Bubble plot showing ligand expression in hDSC (left) and receptor expression in aHFSC (right) across age groups. Color intensity indicates average gene expression, and dot size represents the percentage of positive cells. **k,** Bubble plot showing ligand expression in MACRO (top) and receptor expression in hDSC (bottom) across age groups. Color intensity indicates average gene expression, and dot size represents the percentage of positive cells.

To further explore age-related molecular changes, we analyzed the differentially expressed genes (DEGs) between young (12-20 years old) and old (60-77 years old) individuals, focusing on cells within the hair follicle’s epithelial and mesenchymal compartments. This analysis revealed that hDSC exhibited the highest number of DEGs (2,361 DEGs, *P* < 0.05, |log_2_ fold change| > 0.5) **(Fig. 3c and Supplementary Table 7)**, indicating they are the most transcriptionally dynamic and susceptible cell population during hair follicle aging. Consistent with their stable proportions, both qHFSC and aHFSC showed minimal changes in gene expression level, mirroring observations in mouse model, where aged HFSC maintain their core identity **(Fig. 3c)**^51^. Functional enrichment analysis of hDSC DEGs highlighted several age-associated transcriptional signatures **(Fig. 3d-e, Extended Data Fig. 8b and Supplementary Table 8)**. We observed an upregulation of genes related to oxidative phosphorylation (*NDUFS7, UQCRH, COX6C*) concurrent with a downregulation of glycolysis genes (*HK1, LDHA*), suggesting a potential metabolic shift in aged hDSC. Furthermore, our analysis indicated an enrichment of genes associated with cellular stress, including the unfolded protein response (*DNAJC4, YOD1, EIF2AK3*), oxidative stress (*RELB, FOSL2*), pro-inflammatory signaling (*IL6R, TNFRSF1B, CXCR4*), and cell cycle (*CCND1, CDK2, TOP2A*), alongside a predicted reduction in Hedgehog pathway activity (*PTCH1* high, *GLI2/3* low).

Our multi-omics analysis revealed a robust epithelialization program in aged hDSC, characterized by the upregulation of markers such as *CDH1*, *GRHL3*, and *CLDN1* **(Fig. 3d-e)**^52^. We validated this phenotypic shift *in situ* using immunofluorescence for E-cadherin (CDH1), confirming aberrant epithelialization of aged hDSC **(Fig. 3f)**. Concurrently, cell-adhesion and extracellular matrix genes (*ITGA1*, *ITGA8*, *NPNT*) were downregulated **(Fig. 3e)**, consistent with spatial mapping and *in situ* analyses showing increased hDSC dispersion and reduced abundance in older follicles **(Extended Data Fig. 8c-f).** These observations support a model in which aging hDSC physically detach from the dermal niche.^23^ We also observed a consistent age-dependent decline in senescence-associated secretory phenotype (SASP)^53^ and aging-related gene scores^54^ specifically within hDSC, whereas these signatures increased in epithelial compartments, particularly in TAC populations **(Fig. 3g and Extended Data Fig. 8g-h)**.

### Age-related dynamics of mesenchymal-epithelial stem cells communication

Transcriptomic analysis revealed a marked age-related downregulation of secretion-associated genes in hDSC **(Fig. 3d)**, prompting us to examine potential disruptions in intercellular communication. Leveraging our high-resolution Stereo-seq data, we mapped the physical neighborhoods of hDSC, identifying populations with significant co-localization, including epithelial subsets (TAC1, TAC2, aHFSC), mesenchymal partners (DP, DS), and resident immune cells (macrophages, ILCs, mast cells). Subsequent CellChat^55^ analysis revealed that hDSC exhibited the highest outgoing signaling strength in young follicles, surpassing even that of DP cells. However, this signaling capacity declined with age **(Fig. 3h and Supplementary Table 9)**, suggesting that hDSC-derived signalings plays a dominant role in hair follicle aging.

This global reduction in signaling capacity was driven by a profound, age-dependent loss of key regenerative cues **(Fig. 3i-j and Extended Data Fig. 8i-j)**. Specifically, aged hDSC exhibited a significant downregulation of critical, established growth factors across multiple axes, including WNT (*WNT2*), FGF (*FGF7*), IGF (*IGF1, IGF2*), PDGF (*PDGFA, PDGFD*), and TGFβ (*TGFB2, INHBA*), as well as *GAS6* and *SPP1*^11,56-58^. In parallel, potentially novel hDSC-derived regenerative factors, such as SEMA family members (*SEMA3C, SEMA5A*) and *PTN*, were also downregulated. Compounding this ligand deficit in the mesenchymal niche, we observed a simultaneous downregulation of their cognate receptors within the spatially adjacent epithelial compartments **(Fig. 3j)**. Specifically, receptors for INHBA (*ACVR1B, ACVR2A*), WNT (*LGR5, LRP5, LRP6*), SPP1 (*CD44*), and PTN (*PTPRZ1*) were markedly reduced within the aHFSC population. Similarly, receptors for SEMA3C (*NRP1, NRP2*) were downregulated within the adjacent TACs. This suggests a coordinated mesenchymal-epithelial axis in the aged niche.

Concomitant with the loss of regenerative signals, we observed increased activity of inflammatory interactions (e.g., TIGIT, TWEAK) **(Fig. 3i and Extended Data Fig. 8i)**. Consistent with this pro-inflammatory shift, we identified a progressive, age-dependent upregulation of inflammatory ligands (*IL1A, IL1B, TNF*) originating from the macrophages **(Fig. 3i,k)**. Although the expression levels of the corresponding receptors (*IL1R1, TNFRSF1A*) remained relatively stable in hDSC and HFSC populations **(Fig. 3k and Extended Data Fig. 8k)**, the elevated ligand availability suggests a heightened inflammatory signaling flux in the aged niche. Collectively, these alterations characterized by the erosion of hDSC-derived regenerative signals and the influx of macrophage-driven inflammation likely drive hair follicle miniaturization and impaired regeneration during aging^59^.

### Early hDSC alterations are associated with progressive HFSC dysfunction in AGA

AGA is the most common type of hair loss and features progressive miniaturization of hair follicles^22,60,61^. We integrated the data from 5 AGA patients and 10 age-matched (18-59 years) healthy volunteers to dissect the spatiotemporal dynamics of AGA at single-cell resolution. As expected, hair follicles in AGA were smaller than those in healthy controls, and this miniaturization being particularly pronounced in AF **(Fig. 4a)**. In addition, the proportion of hair follicles in telogen phase was higher in AF, whereas no difference between AO and HO **(Fig. 4b)**. This consistent with the established understanding that AGA disrupts the hair growth cycle, leading to a shortened anagen phase and a prolonged telogen phase^22^.

**Fig. 4.**
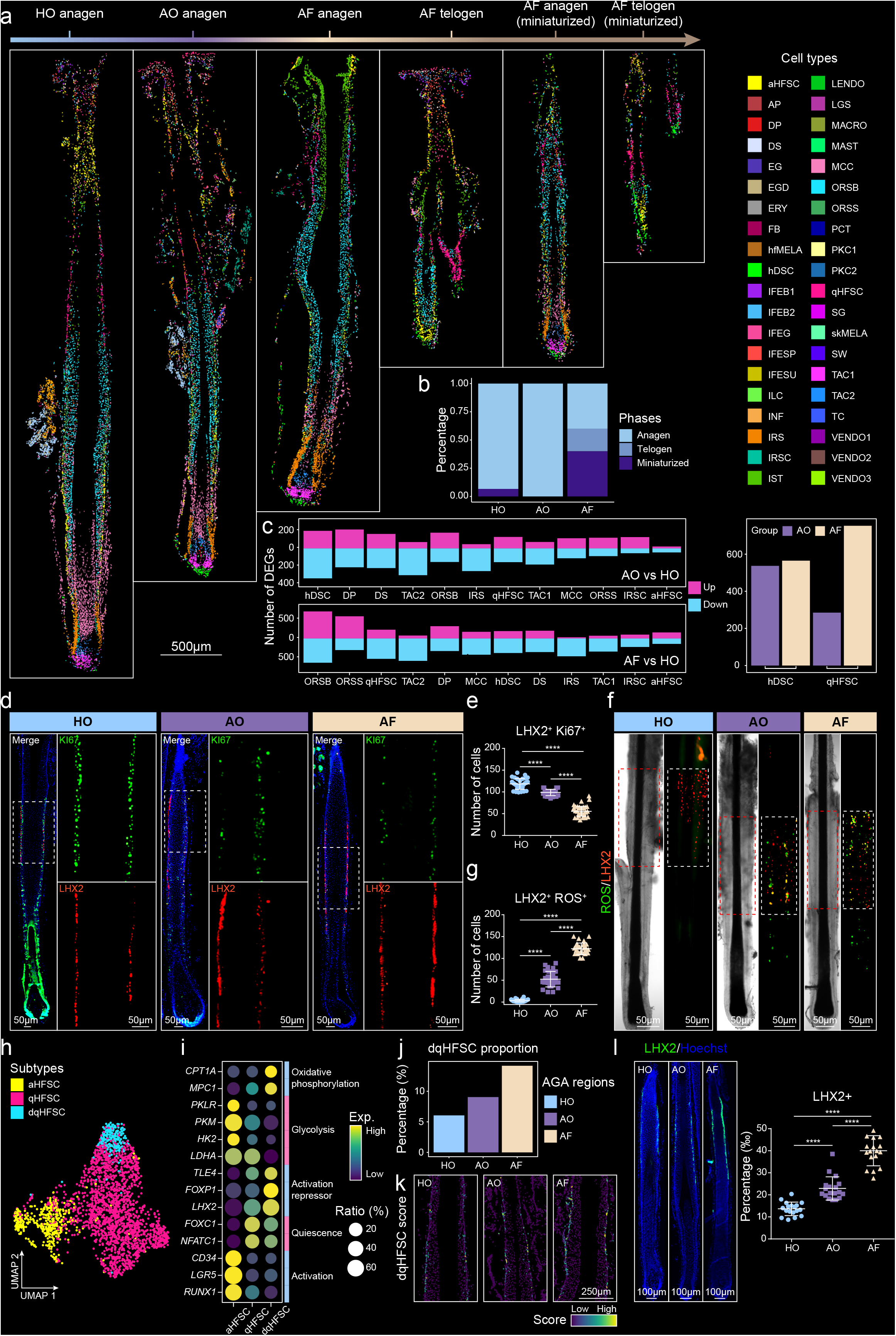
Spatiotemporal cellular state dynamics during AGA. **a,** Spatial visualization of cell type distributions in 6 representative Stereo-seq sections from healthy and AGA samples deconvoluted by RCTD and colored by the 40 identified cell types. HO: healthy occipital scalp; AO: AGA occipital hair-bearing scalp; AF: AGA frontal hairless scalp. **b,** Quantification of the proportion of hair follicles in different states (anagen, telogen, miniaturized) in each group. **c,** Bar plots showing the counts of upregulated (purple) and downregulated (blue) genes comparing AO vs. HO (top) and AF vs. HO (bottom). The right panels summarize the total DEG abundance for hDSC and qHFSC across these comparisons. **d,** Representative immunofluorescence images of hair follicle sections from AGA groups co-staining for Ki67 (green) and LHX2 (red). Scale bar, 50 μm. **e,** Statistical quantification of LHX2^+^Ki67^+^ HFSC populations. Top: Quantification of LHX2^+^Ki67^+^ HFSC in each group (HO, AO, AF). For box plots, the center line denotes the median, and whiskers represent the 1.5 interquartile range. **f,** Representative images of immunofluorescence staining for ROS (green) and LHX2 (red). Scale bar, 50 μm. **g,** Box plot showing the counts of LHX2^+^ ROS HFSC. **h,** UMAP analysis of HFSC subpopulations from scRNA-seq data. Dots are colored by 3 HFSC subtypes: qHFSC, aHFSC, and dqHFSC. **i,** Bubble plot showing the average expression of selected marker genes in the 3 HFSC subtypes. Color intensity indicates average gene expression, and dot size represents the percentage of positive cells. **j,** Quantification of dqHFSC proportions in each group (HO, AO, AF). **k,** Spatial visualization of module scores for dqHFSC marker genes within stage-matched anagen hair follicles from HO, AO, and AF groups. **l,** Representative immunofluorescence staining for LHX2 (green) in human hair follicle sections from AGA groups and box plot showing the proportion of LHX2^+^ HFSC. Scale bar, 100 μm.

To dissect cell-specific changes in different AGA-affected regions, we identified DEGs in various hair follicle cell types by comparing AF and AO to HO **(Fig. 4c)**. We found hair follicle mesenchymal cells, including hDSC, DP and DS cells, exhibited the highest number of DEGs in AO compared to HO. While hair follicle epithelial cells, including ORSB, ORSS cells and qHFSC, displayed the highest number of DEGs in AF compared to HO. These results suggest that AGA initially impacts mesenchymal compartments, even in clinically unaffected occipital regions^62^, with hDSC as primary early responders. Consequently, AO likely represents an early stage of AGA, where initial mesenchymal dysfunction may precipitate subsequent epithelial impairment and more severe hair loss in later stages (AF).

To further explore the transcriptional changes associated with AGA progression, we first focused on hDSC, which exhibited the most pronounced changes in AO **(Extended Data Fig. 9a-d and Supplementary Tables 10-11)**. Functional enrichment analysis of upregulated genes in the AO region revealed an enrichment for apoptosis-associated pathways (*FAS, TNFRSF12A, GADD45A*). In the AF region, we observed increased expression of genes involved in the negative regulation of Wnt signaling (*APCDD1, PRICKLE1, TBX18*). Conversely, genes related to cell cycle progression (*CCNB1, CDC25B, CDK6*) and EGF signaling (*AREG, HBEGF, EGFR*) were downregulated in the AF group. These findings suggest that hDSC exhibit different disease-related transcriptional signatures and play a critical role especially in the early stage of AGA.

Although HFSC dysfunction is thought to contribute to follicle miniaturization during AGA progression, how this process evolves across disease stages remains poorly understood. To dissect this, we performed module score and functional enrichment analysis of DEGs in qHFSC across AGA **(Extended Data Fig. 9e-h and Supplementary Tables 12-13)**. This revealed a progressive metabolic shift: oxidative phosphorylation genes (*NDUFB7, NDUFC2, NDUFA6*) were upregulated from HO to AF, while glycolysis (*HK2, LDHA*) and cell cycle-related programs (*SMC4, YWHAG, UBE2S*) showed a concomitant, stepwise decline (HO > AO > AF). This was accompanied by downregulation of pyruvate dehydrogenase kinases (*PDK1, PDK4*) and increased expression of mitochondrial pyruvate dehydrogenase (*PDHA1*), indicating a gradual uncoupling of glycolytic flux^63,64^. Additionally, protective pathways that sustain stem cell function and glycolysis, including autophagy (*MAP1LC3B, SQSTM1, ATG3*) and the hypoxia response (*HIF1A, SOD2, ATF4*), exhibited a similar progressive decline*^63,65^*.

To validate these transcriptional signatures functionally, we performed immunofluorescence analyses **(Fig. 4d-g)**, revealing a continuous phenotypic gradient: Ki67 expression in HFSC progressively decreased, whereas ROS levels increased from HO to AF. Collectively, these data delineate a continuous pathological trajectory in which qHFSC experience escalating oxidative stress, metabolic reprogramming, and cell cycle arrest. The intermediate phenotype in AO confirms that these molecular perturbations initiate early in clinically non-balding scalp, ultimately driving progressive HFSC dysfunction during AGA.

### Progressive enrichment of deep-quiescence-like HFSC during AGA progression

Given the profound transcriptional and metabolic alterations observed in HFSC during AGA, we sought to determine if these changes correspond to a distinct cellular state. To explore this, we re-clustered the total HFSC pool comprising both qHFSC and aHFSC **(Fig. 4h)**. Beyond the two canonical subpopulations characterized by either activation markers (*LGR5, CD34, RUNX1*)^10^ or quiescence factors (*FOXC1, NFATC1, S100A4*)^34^, we identified a previously uncharacterized cluster whose proportion increased progressively from HO to AF samples **(Fig. 4i-j)**. Notably, this expansion occurred independently of the hair cycle stage **(Fig. 4k and Extended Data Fig. 9i)**. This novel population preserves the core quiescence program (*NFATC1* and *FOXC1*), but selectively accumulates a suite of potent HFSC activation repressors, including *LHX2*, *FOXP1*, and *TLE4* **(Fig. 4i)**^32,43^. We rigorously validated this hyper-repressed signature *in situ* using immunofluorescence, confirming that LHX2 protein levels undergo a continuous, disease-stage-dependent upregulation from HO to AF follicles **(Fig. 4l)**. Functional enrichment analysis further revealed that this state is defined by a signature of cellular exhaustion, characterized by elevated reactive oxygen species (ROS) levels, pro-apoptotic signaling, and cell-cycle arrest **(Extended Data Fig. 9j and Supplementary Table 14)**. Based on this unique molecular profile, in which the superimposition of multiple inhibitory factors likely enforces an insurmountable activation threshold, we designated this population as deep-quiescence-like state (dqHFSC). Pseudotime trajectory analysis further resolved this pathological divergence, uncovering a clear bifurcation originating from the qHFSC state. One branch progressed toward functional activation, while the other deviated toward the dqHFSC fate **(Extended Data Fig. 9k-n)**. Collectively, these data demonstrate that during AGA progression, a subpopulation of qHFSC is driven into a state of deep quiescence, defined by a functional block in proliferation and heightened oxidative stress.

### hDSC in AGA shift from a pro-regenerative to a hyperactive, inhibitory signaling hub

To delineate the intercellular communication architecture in AGA, we performed CellChat analysis across HO, AO and AF groups, incorporating the physical cellular neighborhood constraints defined by our Stereo-seq data **(Extended Data Fig. 10a and Supplementary Table 15)**. This analysis identified hDSC as the principal predicted signaling hub across all conditions. However, the nature and magnitude of their signaling activity undergo a dramatic, stepwise pathogenic shift during disease progression **(Fig. 5a-c)**. In healthy follicles (HO), the signaling output of hDSC is characterized by robust pro-regenerative cues. These include established factors such as EGF (*TGFA, AREG, HBEGF, EREG*)^66-68^, FGF (*FGF10*)^11^, NRG (*NRG1, NRG2, NRG3*), *WNT10B*^69^, alongside the potentially novel regulators *SEMA3C* and *PTN*. As the disease progresses from HO to AF, these regenerative signals are progressively attenuated. This loss aligns with the high expression of their cognate receptors on aHFSC, including *EGFR* for EGF, *NRP1* and *NRP2* for SEMA3C, and *PTPRZ1* for PTN.

**Fig. 5.**
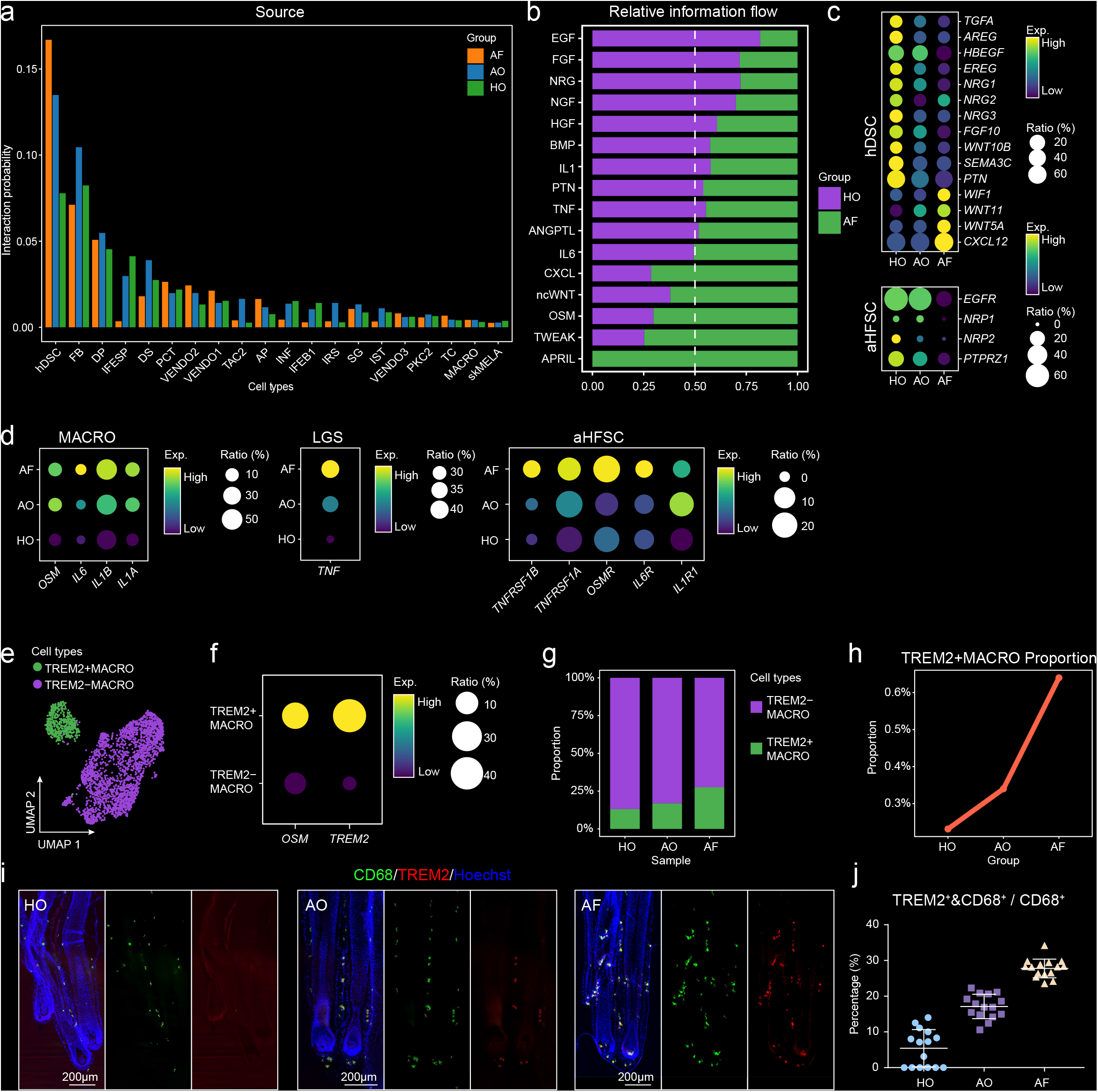
hDSC shift from a pro-regenerative to a hyperactive, inhibitory signaling hub in AGA. **a,** Bar plot showing the signaling interaction strength of each cell type as source (ligand) in HO, AO and AF groups, analyzed by CellChat. **b,** Quantitative plot showing the sum of interaction probability differences (relative information flow) of each signaling pathway between HO and AF groups. **c,** Bubble plot showing ligand expression in hDSC (top) and receptor expression in aHFSC (bottom) across sample groups. Color intensity indicates average gene expression, and dot size represents the percentage of positive cells. **d,** Bubble plot showing DEGs in MACRO (left), LGS (middle) and aHFSC (right) across sample groups. Color intensity indicates average gene expression, and dot size represents the percentage of positive cells. **e,** UMAP showing MACRO subpopulations: TREM2^+^ MACRO and TREM2^-^ MACRO. **f,** Dot plot showing markers of MACRO subpopulations. Color intensity indicates average gene expression, and dot size represents the percentage of positive cells. **g,** Bar plot showing the proportion change of each MACRO subpopulation within the MACRO cell population across different sample groups. **h,** Line plot showing the proportion change of each MACRO subpopulation within all cell populations across different sample groups. **i,** Representative immunofluorescence images of MACRO CD68 (green) and TREM2 (red) co-staining in HO (left), AO (middle), and AF (right). Scale bar, 200 μm. **j,** Box plot showing the proportion of TREM2^+^ MACRO.

Concurrently with this loss of regenerative drive, the total signaling output from hDSC paradoxically increased along the disease trajectory (AF > AO > HO). This surge was driven by the progressive gain of inhibitory and inflammatory pathways, most notably WNT inhibitors (*WIF1, WNT11, WNT5A*) and *CXCL12* **(Fig. 5c)**. Collectively, these findings suggest that hDSC dysfunction in AGA is characterized not simply by the loss of supportive signals, but by the active establishment of an inhibitory microenvironment that progressively suppresses hair follicle regeneration.

### Progressive pro-inflammatory immune remodeling orchestrates regenerative failure in AGA

Given inflammatory enrichment, we specifically interrogated the immune crosstalk within the AGA niche. We identified extensive interaction networks characterized by the progressive upregulation of pro-inflammatory signaling from HO to AF **(Fig. 5d)**. Specifically, macrophages exhibited a stepwise elevation in the expression of *IL1A*, *IL1B*, *IL6*, and *OSM*, while Langerhans cells displayed increased *TNF* expression tracking with AGA stage. This ligand upregulation aligns precisely with the expression of cognate receptors (*IL1R1*, *IL6R*, *OSMR*, *TNFRSF1A*/B) on aHFSC. These findings are consistent with the established roles of IL-1, TNFα and IL6 as potent inhibitors of hair follicle growth that target HFSC in various pathologies, including obesity-induced hair thinning^59^ and AGA pathogenesis^70,71^. For instance, it has been reported that obesity induces excess reactive oxygen species (ROS) and NF-κB activation within HFSC via autocrine and/or paracrine IL-1R signaling^59^. This mechanism mirrors our observation that high levels of IL-1 producing macrophages are associated with HFSC exhibiting elevated oxidative stress in AGA.

Of particular mechanistic significance, we identified a specific expansion of *TREM2*^+^ macrophages across AGA samples **(Fig. 5e-h)**. Immunofluorescence confirmed a significant enrichment of TREM2^+^ macrophages in AGA tissue, particularly within the lesional frontal scalp, where they reside in close physical proximity to both the hDSC and HFSC niches **(Fig. 5i-j)**. This finding provides a translational link to a recent murine study demonstrating that TREM2^+^ macrophages secrete OSM to actively maintain HFSC quiescence and inhibit hair growth^72^.

Together, these findings support a model in which progressive immune remodeling, characterized by enhanced macrophage-derived inflammatory signaling and accumulation of TREM2^+^ macrophages, accompanies the transition toward a chronic anti-regenerative niche during AGA progression. This inflammatory microenvironment may cooperate with the pathogenic transformation of hDSC to promote HFSC dysfunction and regenerative failure.

### hDSC-derived PTN functions as a conserved regenerative cue

To delineate the relationship between physiological aging and AGA, we systematically compared transcriptional alterations in hDSC and HFSC populations across both conditions. Although only a limited number of genes were coordinately regulated in aging and AGA **(Fig. 6a-d)**, module score analysis revealed that these shared programs underwent markedly greater changes during AGA progression than during physiological aging **(Fig. 6b,d)**, indicating an accelerated deterioration of regenerative capacity in the disease state.

**Fig. 6.**
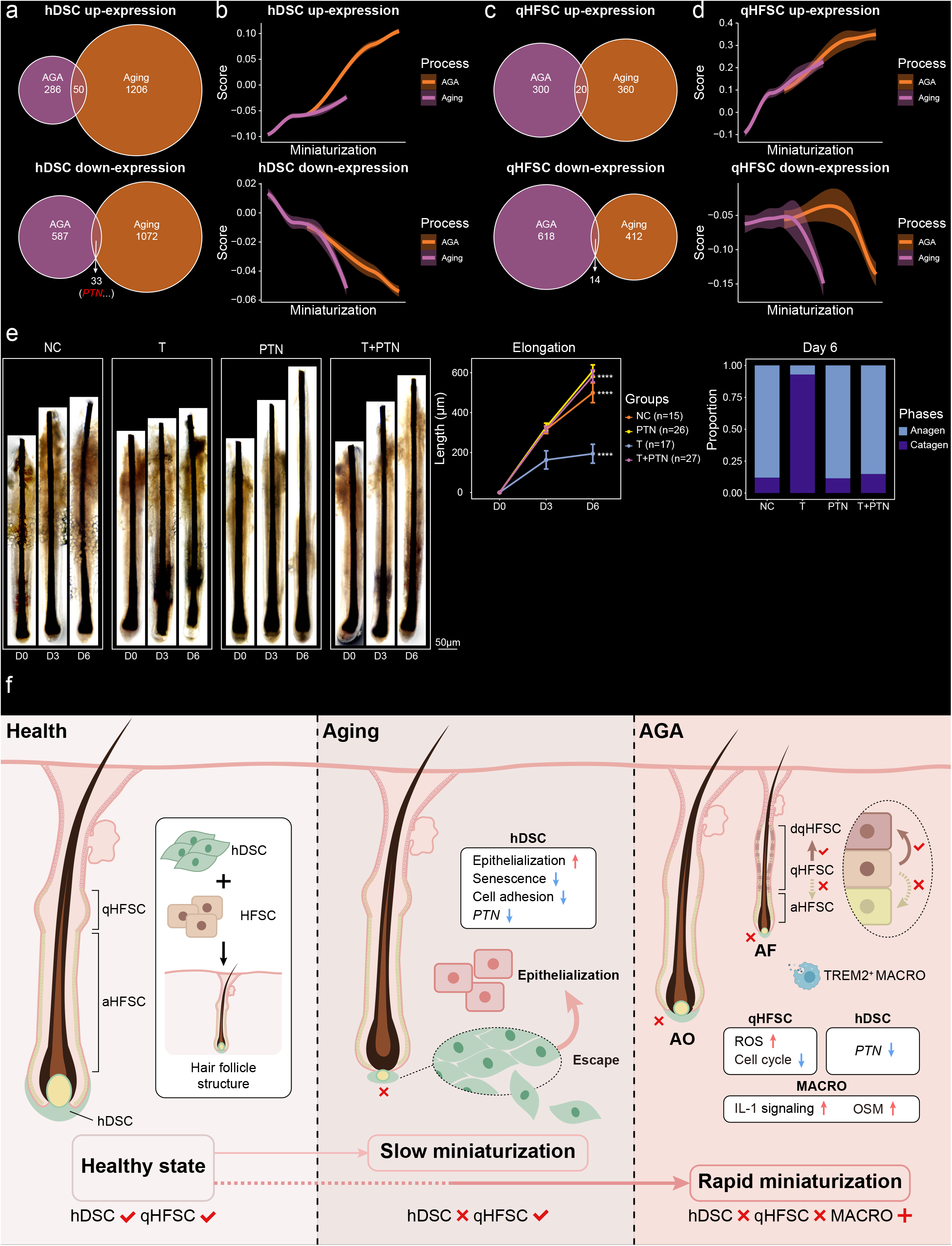
Spatiotemporal events in human hair follicle homeostasis, aging and AGA pathogenesis. **a,** Venn diagrams showing the overlapping co-upregulated (top) and co-downregulated (bottom) gene signatures between aging and AGA within hDSC populations. **b,** Fitting curves illustrating module scores for hDSC co-upregulated (top) and co-downregulated (bottom) genes from (a) against the progression of hair follicle miniaturization. **c,** Venn diagrams showing the overlap of co-upregulated (top) and co-downregulated (bottom) gene sets between aging and AGA within qHFSC populations. **d,** Fitting curves illustrating module scores for qHFSC co-upregulated (top) and co-downregulated (bottom) genes from (c) against the progression of hair follicle miniaturization. **e,** Representative images and quantification of hair shaft elongation and hair follicle cycle staging in *ex vivo* organ-cultured human hair follicles. **f,** The human hair follicle exhibits remarkable regenerative capacity orchestrated by multiple stem cell populations, including HFSC and hDSC. However, both aging and AGA perturb this homeostatic balance, ultimately driving follicular miniaturization and hair loss. Aging is characterized by a numerical decline in the hDSC pool, accompanied by increased spatial dispersion that suggests their progressive detachment from the dermal niche. While aged hDSC undergo a phenotypic transition toward an epithelial-like state, they paradoxically manifest a diminished senescence signature, alongside compromised cell-cell adhesion and downregulated *PTN* expression. AGA initially disrupts the mesenchymal compartment, particularly hDSC, primarily manifested by the downregulated expression of *PTN*. These molecular alterations likely trigger a metabolic shift in qHFSC, promoting their transition into a dysfunctional state (dqHFSC) characterized by ROS accumulation and suppressed cell-cycle reentry. Furthermore, the AGA microenvironment recruits an expanded population of *TREM2*^+^ macrophages characterized by elevated IL-1 signaling and *OSM* levels, which collectively reinforce HFSC quiescence.

From the subset of factors significantly downregulated in both aging and AGA, we identified PTN as a high-priority candidate for functional validation. To evaluate the pro-regenerative potential of this hDSC-derived factor, we utilized an *ex vivo* human hair follicle organ culture model. Supplementation with recombinant human PTN significantly promoted hair shaft elongation compared to vehicle controls **(Fig. 6e)**. Notably, co-treatment with PTN was sufficient to completely reverse the severe hair growth inhibition induced by androgen treatment in this model. These data provide direct experimental proof that the loss of hDSC-derived PTN represents a critical signaling deficit in both the aging and AGA microenvironments, identifying this specific molecule as a potent target for restoring human hair follicle renewal.

## Discussion

Across diverse mammalian tissues and organs, distinct populations of mesenchymal and epithelial stem cells operate to maintain and regenerate their respective lineages^73^. Although these two stem cell compartments frequently coexist within regenerative tissues, the mechanisms through which they communicate to sustain long-term organ renewal remain incompletely understood. The hair follicle provides an ideal model for addressing this question because it contains both epithelial and mesenchymal stem cell populations that must act in concert to support cyclical regeneration. By integrating single-cell transcriptomics, chromatin accessibility profiling, and spatial transcriptomics across homeostasis, aging, and AGA, our study reveals a previously unrecognized framework in which mesenchymal– epithelial stem cell communication functions as a central regulator of human hair follicle regeneration.

A major finding of this study is the identification and characterization of human hair follicle dermal stem cells (hDSC), a previously underappreciated mesenchymal stem cell population that exhibits substantial divergence from murine dermal compartments. Unlike rodent hair follicles, which undergo rapid and highly synchronized regenerative cycles, human scalp follicles sustain an exceptionally prolonged anagen phase that can last for years. The cellular mechanisms supporting this unique regenerative capacity have remained poorly understood.

Our data identify hDSC as the dominant signaling hub within the human follicular niche. Spatial analyses demonstrate that hDSC reside adjacent to activated HFSC and engage in extensive regenerative communication with epithelial compartments. This intense and localized mesenchymal-epithelial stem cell cross-talk likely provides the critical and sustained molecular engine required to maintain the uniquely prolonged human anagen phase. It establishes that functional mesenchymal-epithelial stem cell cross-talk is an prerequisite for successful hair follicle regeneration. More broadly, such reciprocal communication may represent a general principle guiding tissue regeneration in complex organs.

The spatiotemporal tracking of this mesenchymal-epithelial cross-talk reveals how the stem cell niche collapses during physiological aging and AGA. In both aging and AGA contexts, hDSC remain the central signaling locus but undergo profound functional deterioration. Both aging and AGA are characterized by a marked attenuation of pro-regenerative communication from the hDSC toward aHFSC and transit-amplifying cells. However, AGA exhibits a more aggressive pathogenic transformation where the decline in pro-regenerative signals is accompanied by a surge in active inhibitory signals. These distinct niche alterations converge on the epithelial compartment. During AGA progression, HFSC undergo progressive metabolic reprogramming, oxidative stress accumulation, and cell-cycle suppression. This process culminates in the emergence of a deep-quiescence-like HFSC state (dqHFSC), characterized by retention of core stem cell identity together with accumulation of multiple activation repressors. Our findings therefore suggest that regenerative failure in AGA arises not from stem cell depletion, but from pathological stabilization of a quiescent state that prevents normal activation.

Immune remodeling further contributes to this process. Both aging and AGA exhibit increased inflammatory signaling, particularly through macrophage-derived cytokines. However, AGA is distinguished by the accumulation of TREM2^+^ macrophages that secrete OSM to actively suppress hair follicle regeneration. Together, these observations suggest that regenerative failure in AGA results from the combined effects of pathogenic mesenchymal remodeling and chronic inflammatory niche activation.

Although aging and AGA follow distinct molecular trajectories, our comparative analyses revealed a small subset of shared regenerative programs that are progressively lost in both conditions. Among these, PTN emerged as a particularly compelling candidate due to its strong enrichment within hDSC and its consistent downregulation during both aging and disease progression. Functional validation confirmed that hDSC-derived PTN acts as a critical regenerative regulator. More broadly, PTN illustrates how distinct pathological processes can converge upon a common regenerative signaling deficit. Thus, while aging and AGA are mechanistically distinct, both ultimately impair tissue renewal through disruption of a shared mesenchymal–epithelial communication network.

Ultimately, this study elevates the concept of mesenchymal-epithelial stem cell cross-talk from a localized interaction to a fundamental regulatory paradigm. Deciphering how distinct stem cell lineages mutually sustain each other and coordinate their activities provides a transformative framework not only for treating human hair loss disorders but also for advancing the broader fields of organ biology and regenerative medicine **(Fig. 6f)**.

## Extended Data figure legends

**Extended Data Fig. 1.**
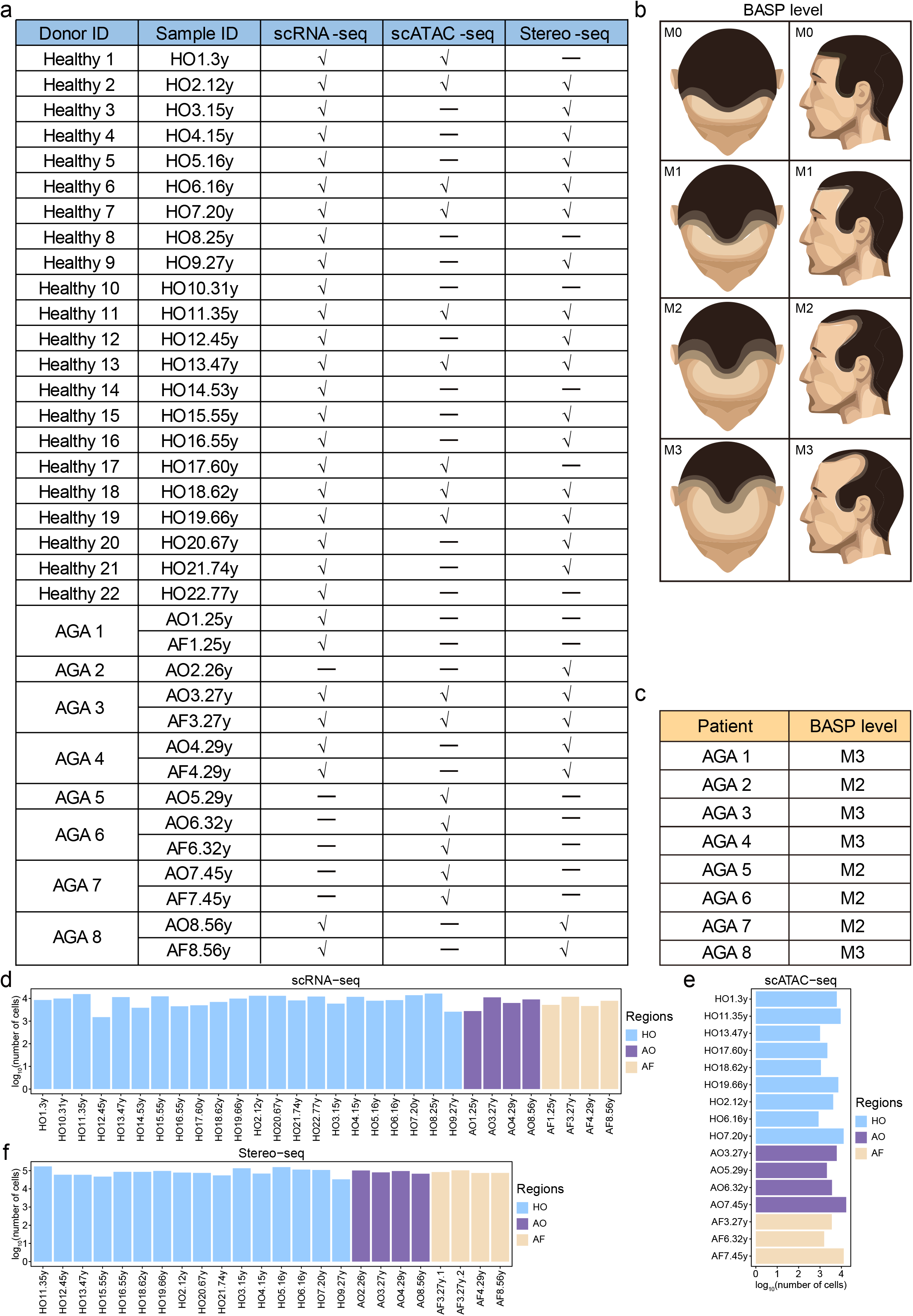
Sample information and AGA severity classification. **a,** List of 30 samples processed for scRNA-seq, scATAC-seq, and Stereo-seq experiments. **b,** Schematic illustration of the diagnostic and grading criteria for AGA from M0 to M3 based on the BASP classification. **c,** The BASP level of each AGA patient included in this study. **d-f,** Quantification of the number of scRNA-seq (d) scATAC-seq (e) and Stereo-seq (f) profiled cells for each sample after quality control.

**Extended Data Fig. 2.**
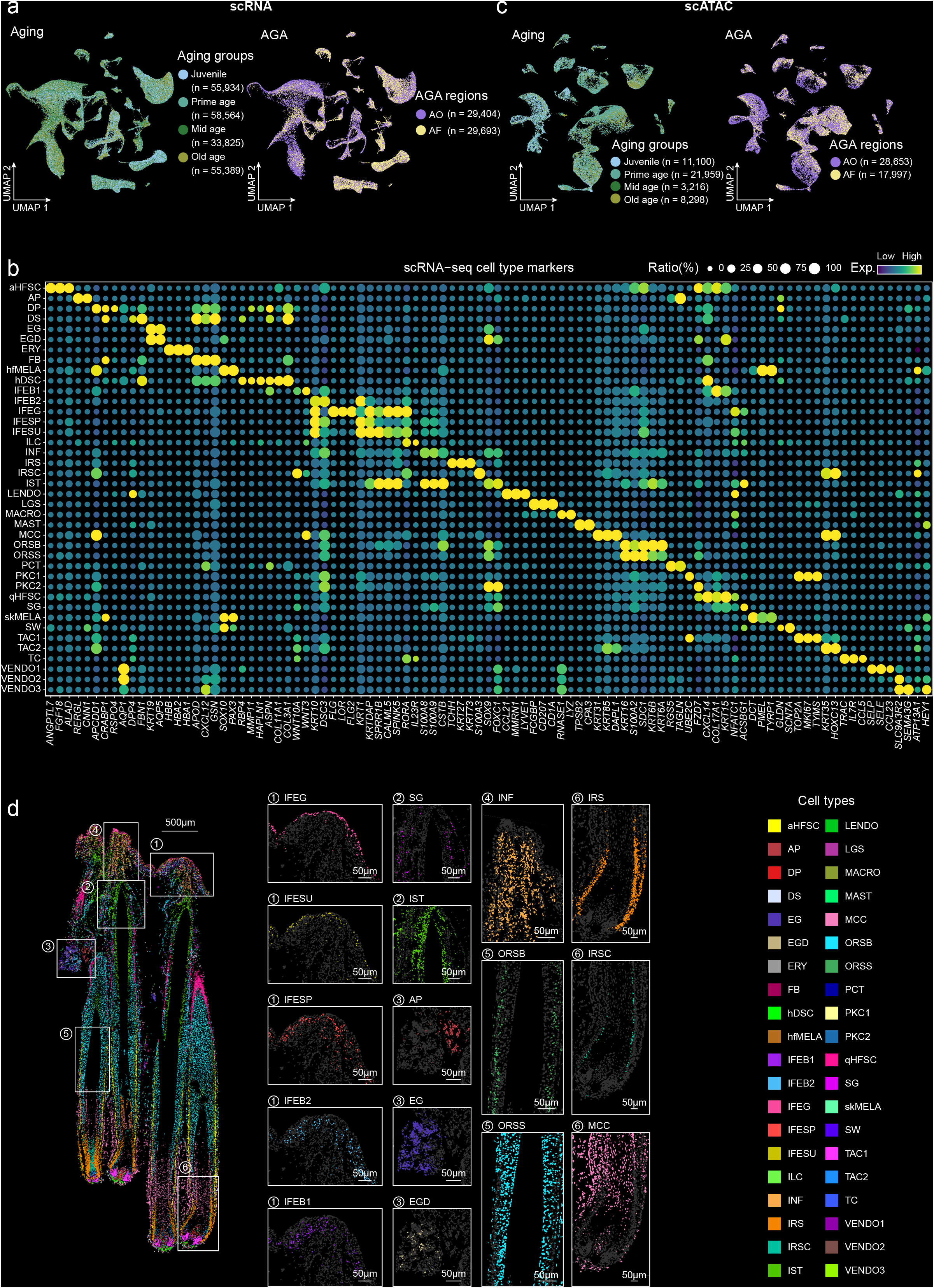
scRNA-seq marker genes and cell spatial characteristics. **a,** UMAP visualization of scRNA-seq profiled cells from aging groups (left) and AGA regions (right). **b,** Bubble plot showing the expression of canonical marker genes for each cell type in scRNA-seq data. Color intensity indicates average gene expression, and dot size represents the percentage of positive cells. **c,** UMAP visualization of scATAC-seq profiled cells from aging groups (left) and AGA regions (right). **d,** Spatial visualization of additional cell type distributions in Fig. 1d. Full view of the hair follicle section, with white boxes indicating magnified areas shown in the right panels (left). Magnified views of selected cell type distributions in the indicated areas (right).

**Extended Data Fig. 3.**
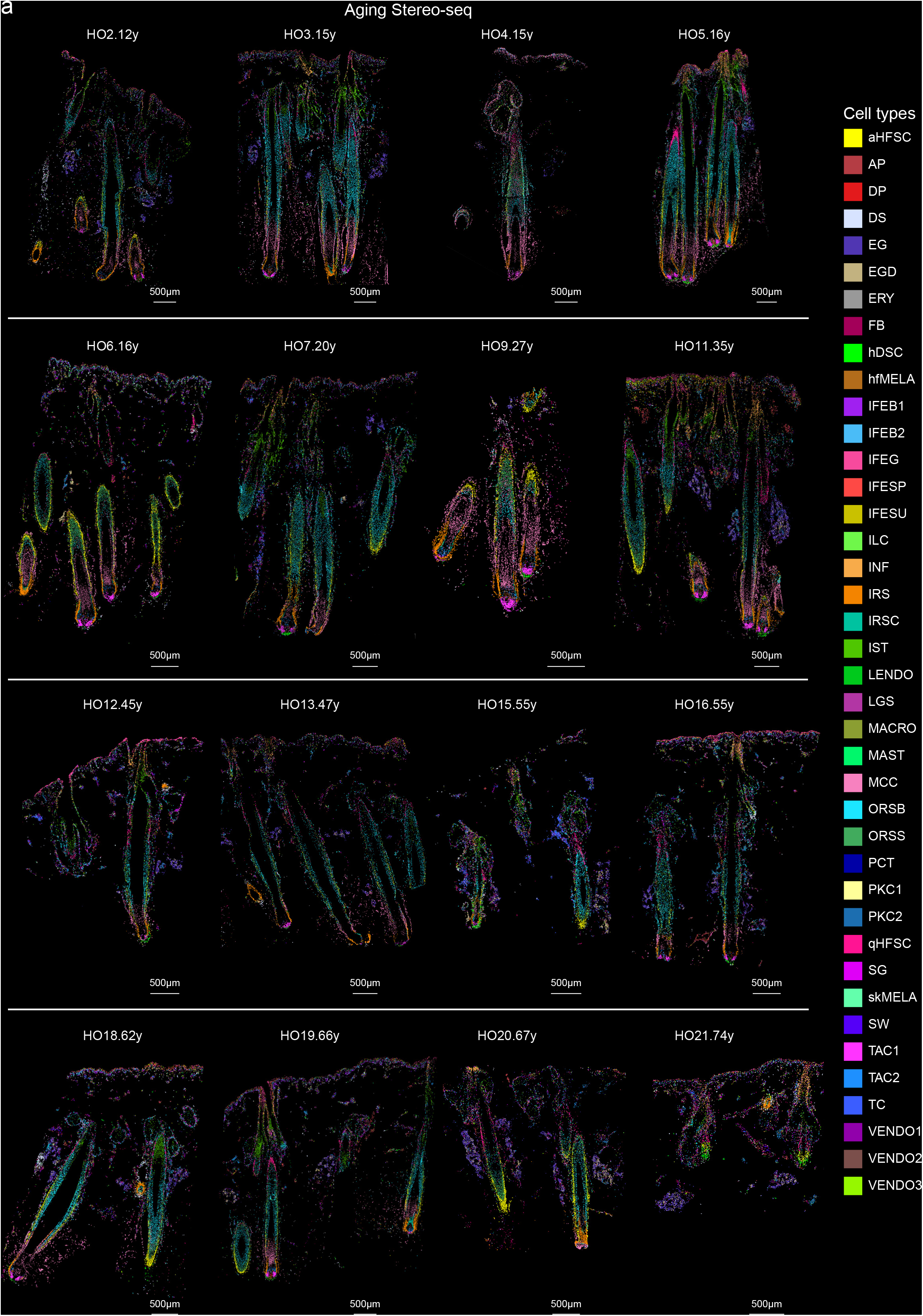
Spatial distribution of cell types in aging human hair follicles. **a,** Spatial visualization of cell type distributions in 16 Stereo-seq sections from aging samples deconvoluted by RCTD and colored by the 40 identified cell types.

**Extended Data Fig. 4.**
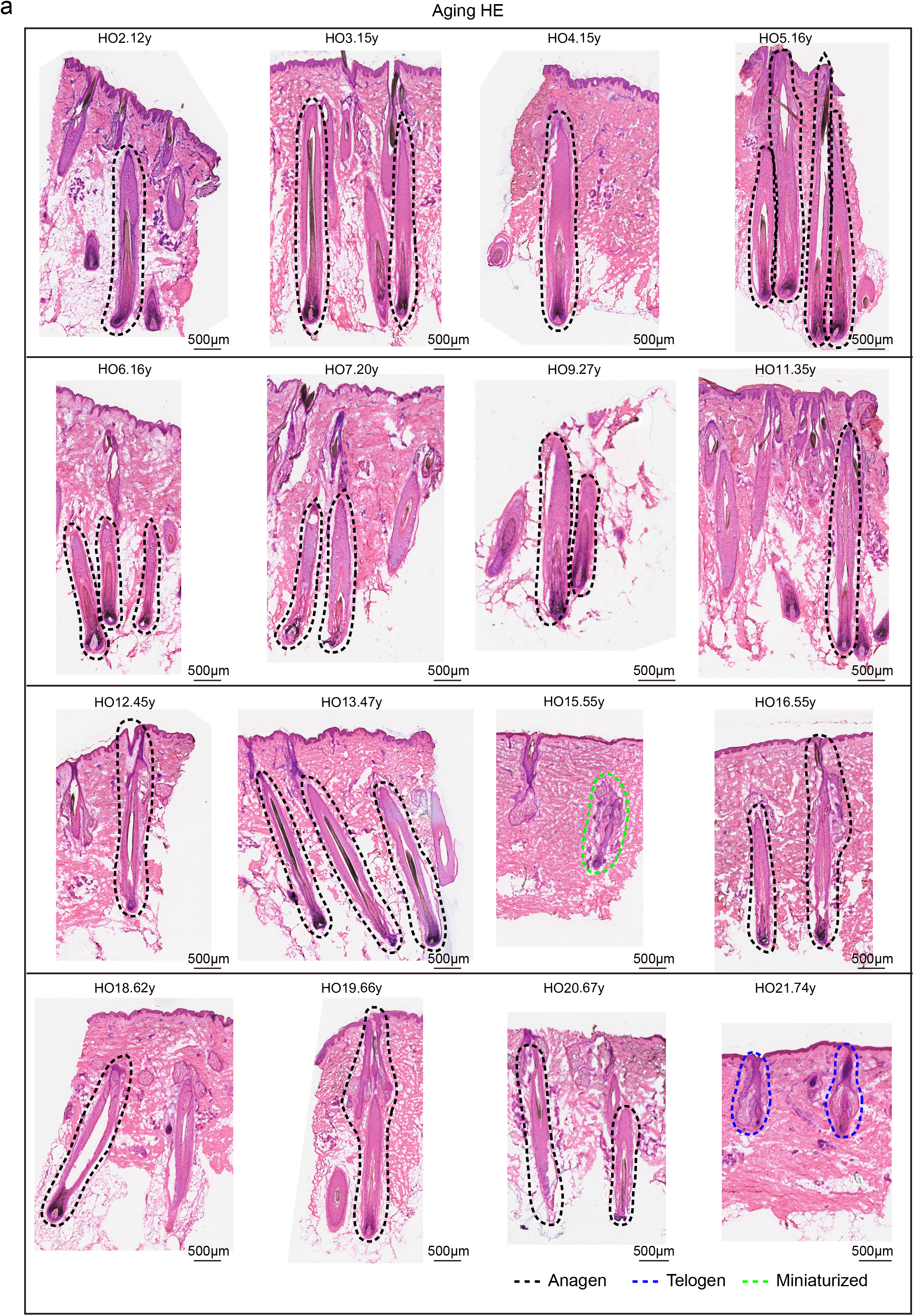
H&E staining of aging samples. **a,** H&E staining of the adjacent sections corresponding to the Stereo-seq sections shown in Extended Data Fig. 3 |Anagen: the hair follicles in anagen phase (black dotted line), telogen: the hair follicles in telogen phase (blue dotted line), miniaturized: the miniaturized hair follicles (green dotted line).

**Extended Data Fig. 5.**
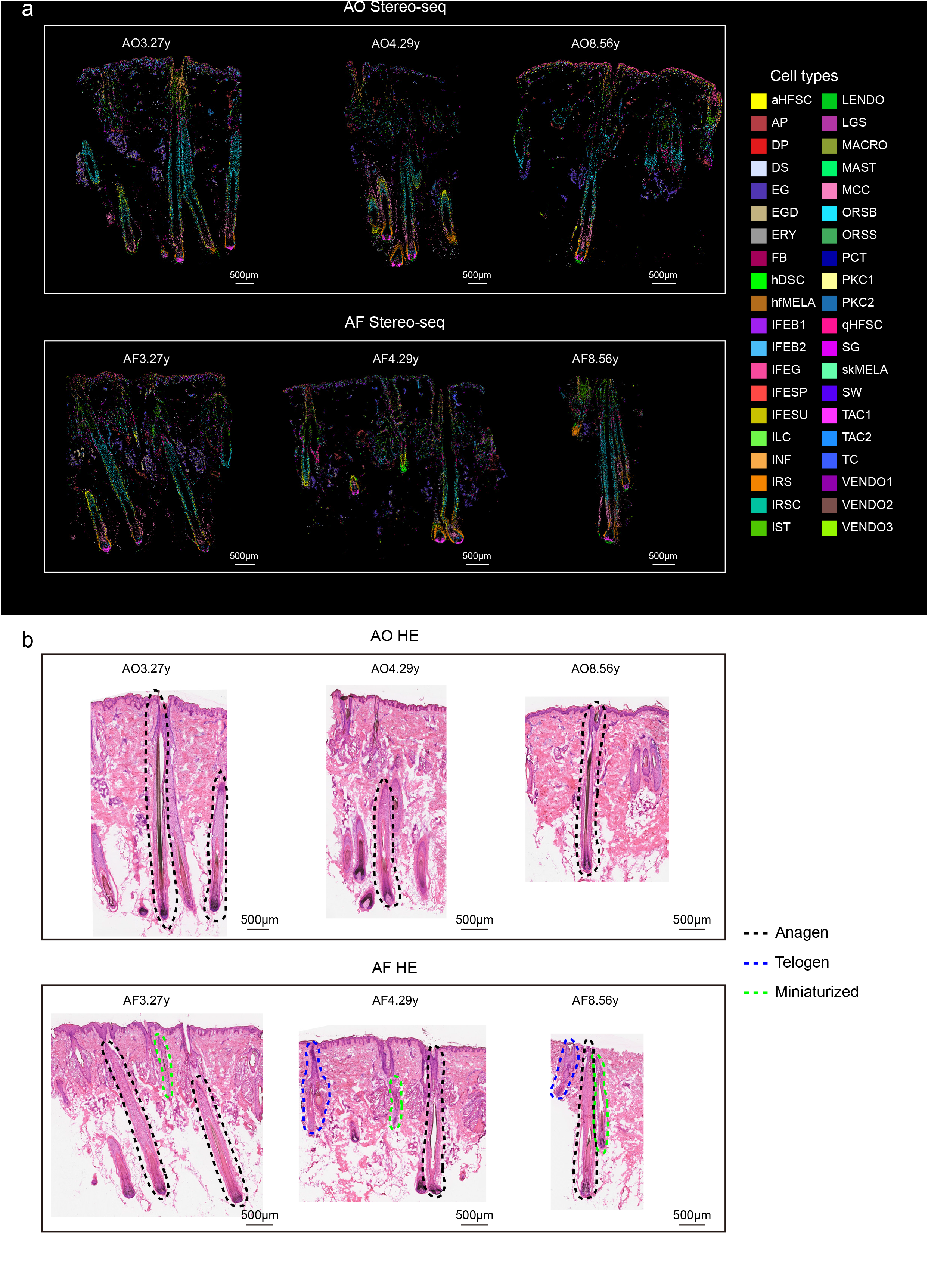
Spatial distribution of cell types in human hair follicles with AGA. **a,** Spatial visualization of cell type distributions in 3 Stereo-seq sections from the AGA occipital regions (top) and frontal regions (bottom) deconvoluted by RCTD and colored by the 40 identified cell types. **b,** H&E staining of the adjacent sections corresponding to the Stereo-seq sections shown in (a). Anagen: the hair follicles in anagen phase (black dotted line), telogen: the hair follicles in telogen phase (blue dotted line), miniaturized: the miniaturized hair follicles (green dotted line).

**Extended Data Fig. 6.**
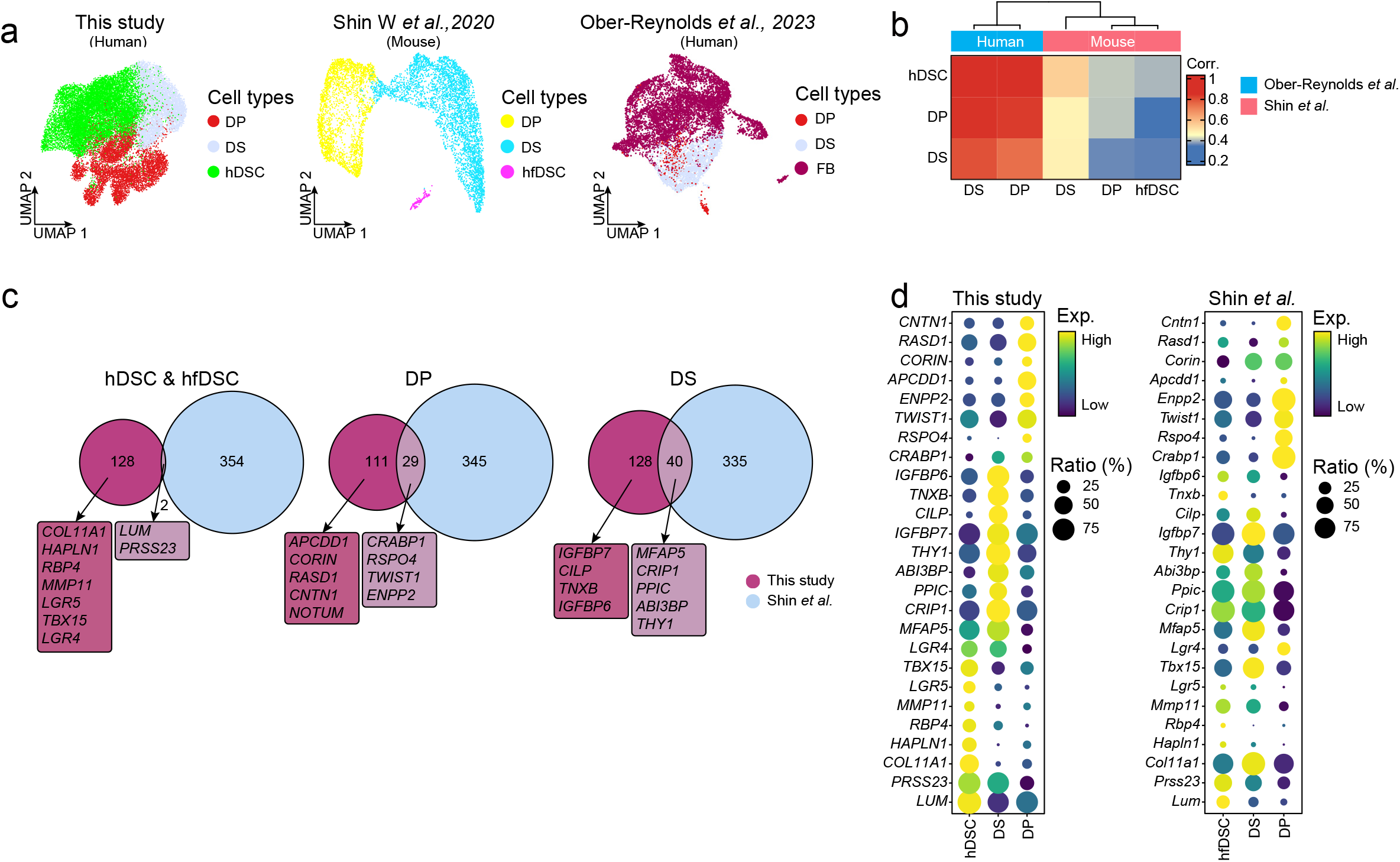
HFSC marker spatial expression and interspecies conservation. **a,** UMAP visualization of human hair follicle dermal cells (DP, DS, and hDSC) from this study (left), mouse hair follicle dermal cells (DP, DS, and hfDSC) taken from the scRNA-seq dataset of Shin *et al*., 2020^13^ (middle) and human hair follicle dermal cells (DP, DS, and FB) taken from the scRNA-seq dataset of Ober-Reynolds *et al*., 2023^28^ (right). **b,** Heatmap of Spearman correlation showing transcriptional conservation of hair follicle mesenchymal lineage across species and datasets (human DS, mouse hfDSC, and our hDSC lineage). **c,** Venn diagram of conserved and unique hair follicle dermal cell signatures between human and mouse. **d,** Bubble plot showing the interspecies comparison of hDSC lineage-specific genes. Color intensity indicates average gene expression, and dot size represents the percentage of positive cells.

**Extended Data Fig. 7.**
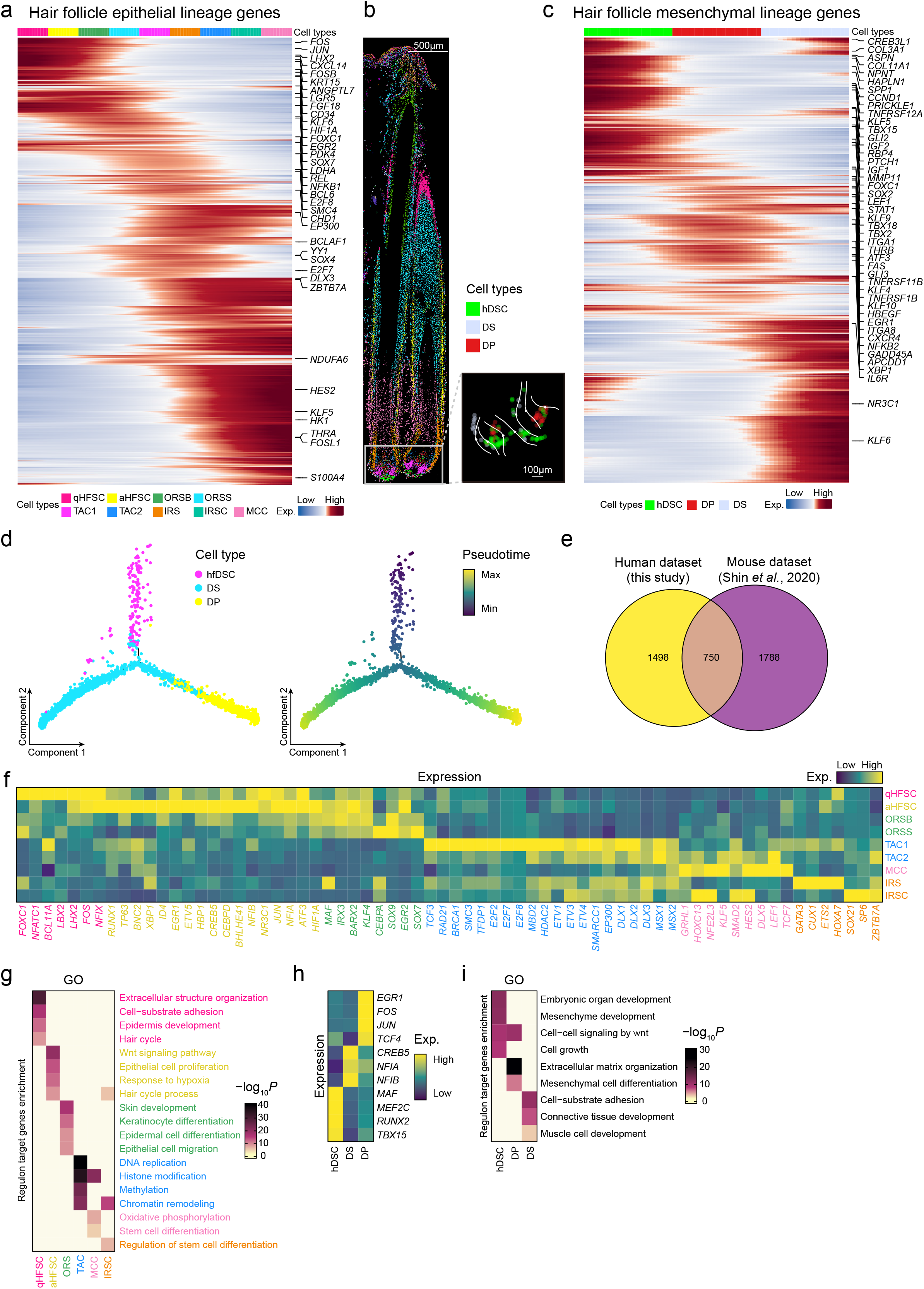
Integrated single-cell transcriptomic and epigenomic landscapes of hair follicle epithelial and mesenchymal cell lineages. **a, c,** Heatmap showing the expression of genes identified along the hair follicle epithelial **(a)** and mesenchymal **(c)** cell lineage trajectories. Colors indicate relative expression levels. The annotations above the heatmap represent cell types. **b,** RNA velocity streamline plot showing the predicted trajectory of hair follicle mesenchymal cell lineages in a representative Stereo-seq section. **d,** Pseudotime trajectory analysis of hfDSC population in the scRNA-seq dataset using Monocle 2. Dots are coloured by mouse hair follicle mesenchymal cell lineages (left) and pseudotime (right). **e,** Venn diagram showing the overlap of hair follicle mesenchymal lineage trajectory genes between human (current study) and mouse (Shin *et al.*). **f,h,** Heatmap showing the average expression of identified regulons across the hair follicle epithelial **(f)** and mesenchymal cell types **(h)**. **g, i,** Heatmap of functional enrichment analysis of target genes corresponding to the hair follicle epithelial (g) and mesenchymal (i) lineage-specific regulons. Color intensity represents pathway significance (-Log_10_ *P*).

**Extended Data Fig. 8.**
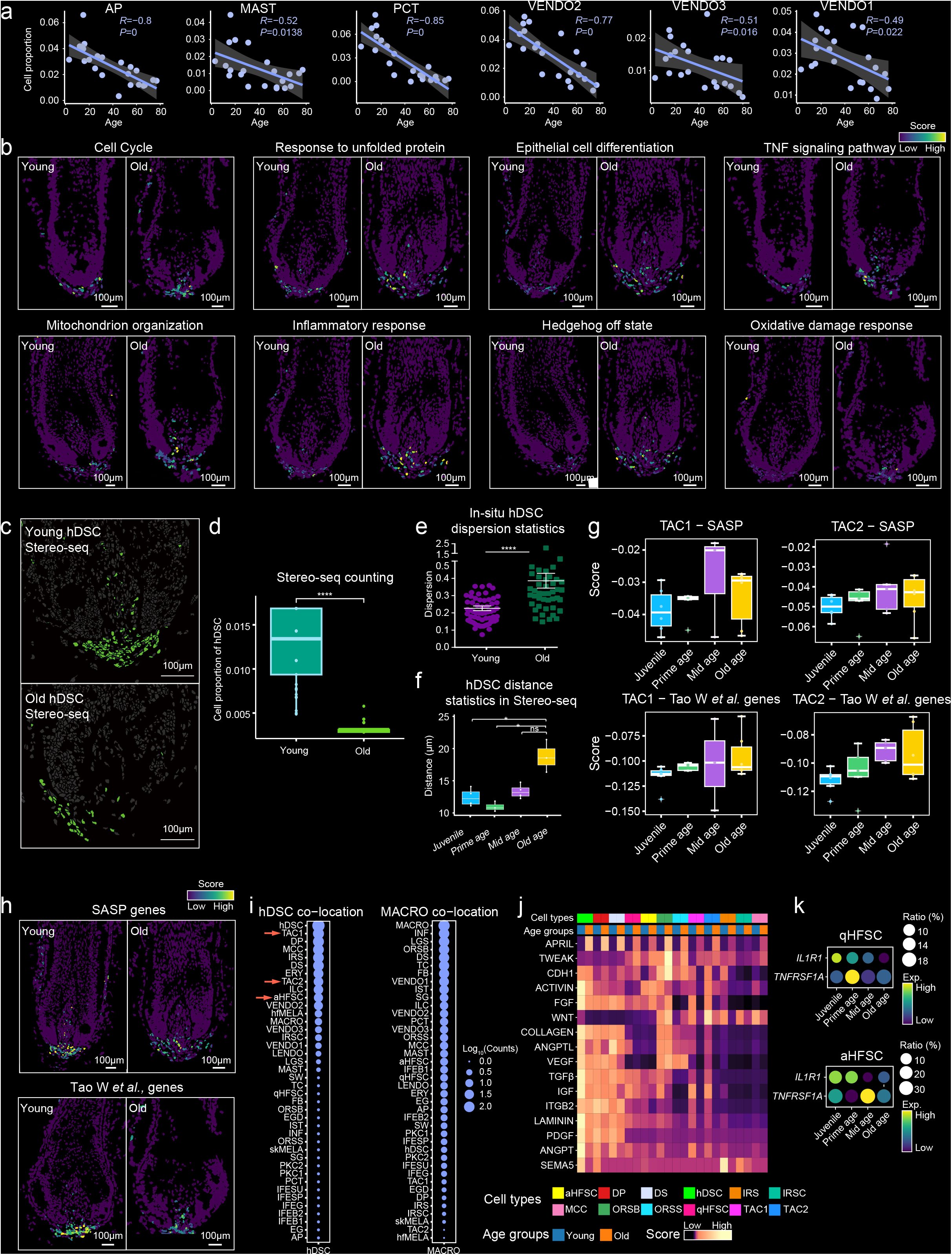
Cell proportion and interaction in aging hair follicles. **a,** Correlation plots showing the relationship between the proportion of indicated cell types and age. *R*, Pearson correlation coefficient; *P*, two-sided *P*-value; the center represents the average value; shading indicates the 95% confidence interval. **b,** Spatial visualization of gene modules for DEGs associated with indicated biological process from Fig. 3d in young (left) and old (right) Stereo-seq sections. **c,** Spatial visualization of hDSC distribution in representative Stereo-seq sections from young (top) and old (bottom) groups. **d,** Box plot showing the proportion of hDSC in young and old groups based on Stereo-seq data. **e,** Quantification of the degree of spatial distance dispersion for hDSC based on immunofluorescence staining from Fig. 3f. **f,** Box plots showing hDSC spatial distance dispersion from Stereo-seq sections across indicated age groups. Boxes denote upper/lower quartiles, lines indicate the median, and whiskers show 1.5-fold interquartile range. *P*-values were determined by two-tailed Student’s t-test. **g,** Box plots showing the module scores of senescence-associated secretory phenotype (SASP) genes (top) and aging-related genes taken from Tao *W et al.*, 2024 (bottom) in TAC1 and TAC2 from indicated age groups. **h,** Spatial visualization of gene modules for SASP genes (top) and age-related genes taken from Tao *et al.*, 2024^54^ study (bottom) in young (left) and old (right) Stereo-seq sections. **i,** Bubble plot illustrating average cell-cell colocalization scores in Stereo-seq sections from aging groups. Dot size represents the Log_10_ (number of colocalized cells). **j,** Heatmap showing the strength of interaction probability for indicated signaling pathways in each cell types between young and old groups. **k,** Bubble plot showing receptor expression in qHFSC (top) and aHFSC (bottom) across age groups. Color intensity indicates average gene expression, and dot size represents the percentage of positive cells.

**Extended Data Fig. 9.**
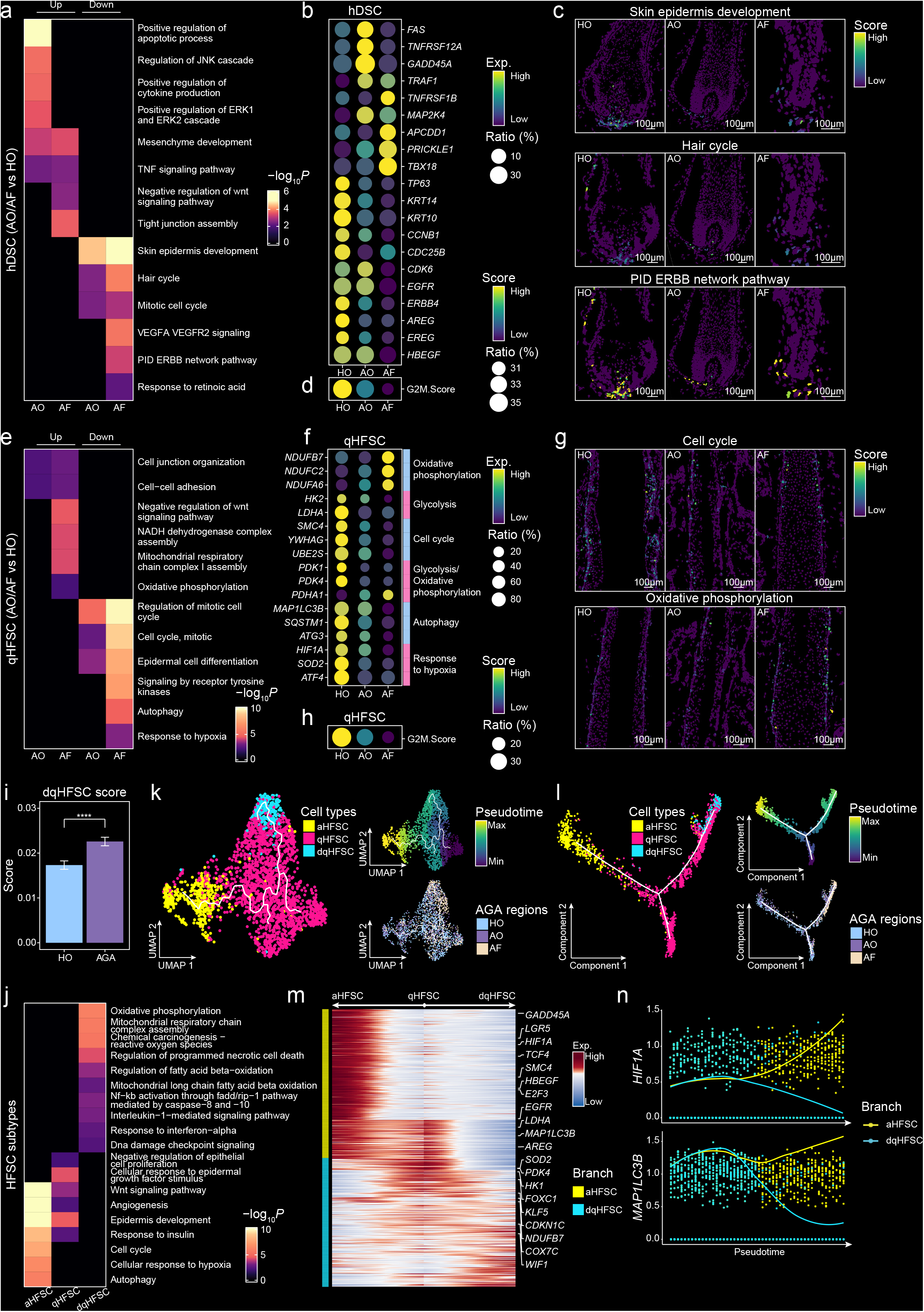
Molecular changes in HFSC and hDSC during AGA. **a,** Heatmap of functional enrichment analysis of upregulated and downregulated DEGs in hDSC comparing AO and AF to HO. Color intensity represents pathway significance (-Log_10_ *P*). **b,** Bubble plot showing the average expression of selected hDSC DEGs related to the indicated biological processes in different groups (HO, AO, AF). Color intensity indicates average gene expression, and dot size represents the percentage of positive cells. **c,** Spatial visualization of gene modules for DEGs associated with indicated biological process from **(a)** in Stereo-seq sections from different groups (HO, AO and AF). **d,** Bubble plot showing the G2M score for hDSC in different groups. Color intensity indicates average gene expression, and dot size represents the percentage of positive cells. **e,** Heatmap of functional enrichment analysis of upregulated and downregulated DEGs in qHFSC. Color intensity represents pathway significance (-Log_10_ *P*). **f,** Bubble plot showing the average expression of selected qHFSC DEGs related to the indicated biological processes in different groups (HO, AO, AF). Color intensity indicates average gene expression, and dot size represents the percentage of positive cells. **g,** Spatial visualization of gene modules for DEGs associated with indicated biological process from **(e)** in Stereo-seq sections from different groups (HO, AO and AF). **h,** Bubble plot showing the G2M score for qHFSC in different groups. Color intensity indicates average gene expression, and dot size represents the percentage of positive cells. **i,** Bar plot comparing dqHFSC signature scores in anagen hair follicles from healthy controls (HO) and AGA Stereo-seq samples. \*\*\*\**P* < 0.0001 (Wilcoxon rank-sum test/Student’s t-test). **j,** Heatmap of functional enrichment analysis of marker genes in each HFSC subtype compared to the other subtypes. Color intensity represents pathway significance (-Log_10_ *P*). **k,** Pseudotime trajectory analysis of HFSC population in the scRNA-seq dataset using Monocle 3. Dots are coloured by 3 HFSC subtypes (left), pseudotime (top right) and 3 groups (bottom right). **l,** Pseudotime trajectory analysis of HFSC population in the scRNA-seq dataset using Monocle 2. Dots are coloured by 3 HFSC subtypes (left), pseudotime (top right) and 3 groups (bottom right). **m,** Heatmap showing the expression of genes identified along the HFSC state trajectory using Monocle 2. Colors indicate relative expression levels. **n,** Expression values of *HIF1A* (top) and *MAP1LC3B* (bottom) across pseudotime. The x-axis represents pseudotime values learned for each cell, and the y-axis represents z-scored Log_2_ (gene expression values).

**Extended Data Fig. 10.**
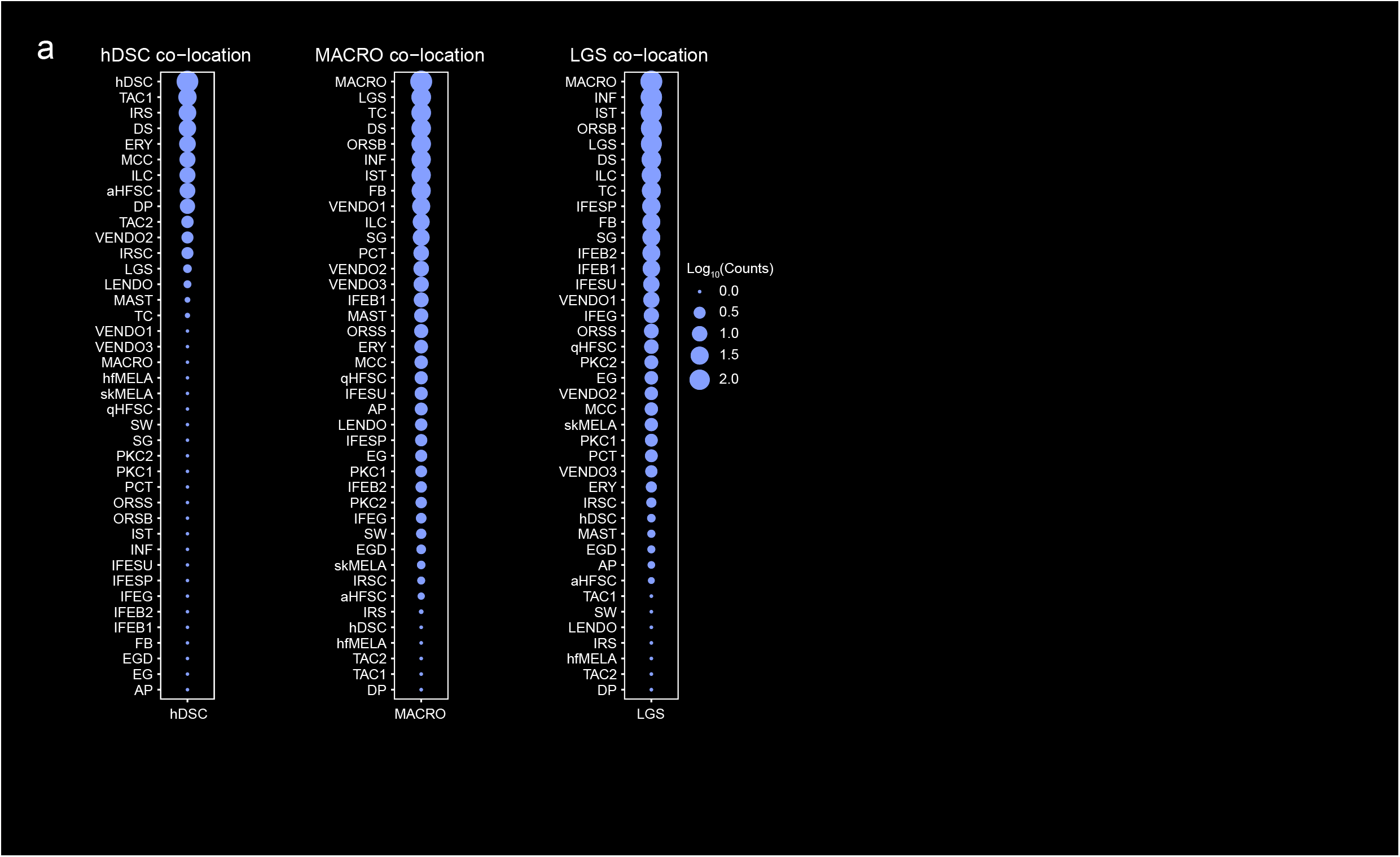
Immune crosstalk in AGA. **a,** Bubble plot illustrating average cell-cell colocalization scores in Stereo-seq sections from AGA groups. Dot size represents the Log_10_ (number of colocalized cells).

## Methods

### Study participants and experimental models AGA diagnosis and ethical clearance

Human scalp samples were collected with informed consent from 22 male healthy volunteers undergoing scalp tumor excision surgery, and 6 male AGA patients (BASP classification 2-3) of non-balding occipital and balding frontal sites undergoing hair transplantation surgery. For 6 individuals below 18 years, the informed consent was obtained from the legally acceptable representative. The diagnosis of AGA, as well as the determination of grading, was determined after an interview by two hair specialist doctors. Ethical approval was granted by the Ethics Committee of Hangzhou First People’s Hospital (ZN-20230617-0124-01).

### Experimental procedures Single cell isolation

Single-cell suspensions were prepared according to a previously published protocol^74^. Briefly, human scalp tissues were rinsed with cold phosphate-buffered saline (PBS) to remove debris, and subcutaneous fat was carefully removed using a sterile scalpel under a dissecting microscope. Tissues were then minced into small pieces using sterile ophthalmic scissors. Minced tissues were incubated in 5 mL of prewarmed Enzyme I solution (4.0 mg/mL dispase II) at 37°C for 50 minutes, followed by prewarmed Enzyme II solution (1.6 mg/mL collagenase type I and 2.0 mg/mL collagenase type IV) at 37°C for 30 minutes. The digestion was stopped by adding 20 mL of Dulbecco’s Modified Eagle Medium (DMEM), and the cell suspension was passed through a 70 μm cell strainer. The collected filtrate was set aside, and the remaining undigested tissue fragments were further digested with 8 mL of 0.25% trypsin at 37°C for 15 minutes. This second digestion was halted by adding 10 mL of DMEM containing 20% fetal bovine serum (FBS). The cell suspension was then filtered through a 70 μm cell strainer, combining the filtrate with the previously collected filtrate. All subsequent procedures were performed on ice or at 4 °C. The pooled cell suspension was centrifuged, and the cell pellet was resuspended in PBS containing 2% FBS. This suspension was then passed through a 30 μm cell strainer to remove any remaining debris. Finally, the filtered suspension was centrifuged, and the cell pellet was resuspended in PBS containing 0.04% bovine serum albumin (BSA) to generate the final single-cell suspension for scRNA-seq and scATAC-seq library preparation.

### scRNA-seq library construction

Libraries for single cell RNA sequencing (scRNA-seq) were prepared following the manufacturer’s instructions for the DNBelab C Series High-throughput Single cell RNA Library Preparation Set V2.5 (MGI, 940-000519-00). Briefly, single cell suspensions were processed for droplet-based cell barcoding, followed by droplet breakage, reverse transcription, second-strand synthesis, and cDNA amplification. Amplified cDNA was then fragmented, ligated to sequencing adapters, and circularized to generate barcoded libraries. All libraries were sequenced on DNBSEQ platform at the CNGB using a paired-end sequencing strategy with the following read lengths: 47 bp for read 1, 100 bp for read 2 and 10 bp for index read.

### scATAC-seq library construction

Libraries for single-cell assay for transposase-accessible chromatin sequencing (scATAC-seq) were prepared according to the manufacturer’s instructions for the DNBelab C Series High-throughput Single-cell ATAC Library Preparation Set V2.0 (MGI, 940-000793-00). Briefly, single-cell nuclei suspensions were subjected to Tn5 transposase tagmentation, followed by droplet-based cell barcoding and transposase reaction in droplets. After droplet breakage, tagged DNA fragments underwent further amplification and were circularized to generate barcoded libraries. All libraries were sequenced on DNBSEQ platform at the CNGB using a paired-end sequencing strategy with the following read lengths: 115 bp for read 1, 69 bp for read 2 and 10 bp for index read.

### Stereo-seq library construction

Fresh scalp tissue samples were washed with PBS, and excess subcutaneous fat was removed. Tissue samples were oriented to preserve hair growth direction and embedded in Tissue-Tek OCT (Sakura, 4583) on dry ice. Embedded tissues were stored at -80°C until further processing.

Libraries for Stereo-seq were prepared according to the manufacturer’s instructions for the Stereo-seq Transcriptomics Set for Chip-on-a-slide (MGI, 201ST114). Briefly, frozen tissue blocks were equilibrated to -20°C and sectioned at 12 μm thickness using a freezing microtome (DAKEWE, CT520). Tissue sections were mounted onto Stereo-seq chips, incubated at 37°C for 5 minutes, and fixed in pre-chilled (-20°C) methanol (Sigma-Aldrich, 34860) for 30 minutes. Sections were then stained with 100 μL of ssDNA fluorescent staining solution for 5 minutes at room temperature in the dark, followed by washing with 0.1× saline-sodium citrate (SSC) buffer and air drying. Glycerol (Solarbio, G8190) was applied to the tissue sections, followed by coverslip placement. Slides were imaged using a fluorescence microscope (Stereo-OR 100, SCI-01-016), and image quality was assessed using ImageQC software. After imaging, coverslips were removed from the chips and washed with 5× SSC buffer (Ambion, AM9770), followed by 0.1% pepsin (Sigma-Aldrich, P7000) permeabilization for 8 min at 37 °C. After permeabilization, in situ reverse transcription was performed on-chip at 42°C for 2 hours. Chips were then washed and transferred to a 24-well plate for tissue removal, which was achieved by incubating chips with tissue removal reagent at 55°C for 10 minutes. Following a wash step, cDNA was released from the chips by incubation with release reagent at 55°C for 3 hours. Released cDNA was amplified by PCR using a ProFlex™ 3 × 32-well PCR System (ABI, 4483636). Amplified cDNA was purified and sequenced on DNBSEQ platform at the CNGB using a paired-end sequencing strategy with the following read lengths: 35 bp for read 1 and 100 bp for read 2.

### Immunofluorescence staining

Human samples were fixed in 4% paraformaldehyde, embedded in paraffin, and sectioned at 7 μm thickness. Paraffin sections underwent deparaffinization and antigen retrieval, followed by blocking with 5% bovine serum albumin (BSA) for 1 hour at room temperature. Sections were then incubated with primary antibodies overnight at 4°C. Following three washes with PBS, sections were incubated with fluorescent-conjugated secondary antibodies for 1 hour at room temperature. Nuclei were counterstained with 10 mg/mL Hoechst 33342 for 15 minutes at room temperature. Images were acquired using an OLYMPUS FV4000 confocal microscope. For each antibody, five to ten random fields were selected for quantification of positively stained cells. The following primary antibodies were used in this study: Mouse anti-E-Cadherin (Abcam, ab231303), Rabbit anti-CK15 (Abcam, ab52816), Rabbit anti-AQP1 (Proteintech, 20333-1-AP), Rabbit anti-SOX2 (Abcam, ab97959), Rabbit anti-DIO2 (Proteintech, 26513-1-AP), Mouse anti-LHX2 (Santa Cruz Biotechnology, sc-374658). Secondary antibodies (Jackson ImmunoResearch) used in this study were Goat Alexa Fluor 488 anti-rabbit IgG (111-545-003) and Goat Alexa Fluor 594 anti-mouse IgG (115-585-003).

### Human hair follicle organ culture

Human terminal hair follicles were obtained from the non-balding occipital scalp of male patients with androgenetic alopecia undergoing elective hair transplantation surgery. Following meticulous microdissection, intact anagen VI-phase follicles were selected and pre-incubated in William’s E medium (ThermoFisher Scientific) for 24 h to allow recovery from isolation stress.

Follicles were then randomly assigned to experimental groups and cultured at 37 °C in a humidified incubator with 5% CO□, using William’s E medium supplemented with 2 mM L-glutamine, 10 ng/mL hydrocortisone, 10 μg/mL recombinant human insulin, and 1% (v/v) penicillin-streptomycin. Follicle growth was monitored every 48 h by stereomicroscopy, with hair shaft elongation measured using image analysis software. Hair cycle stage (anagen, early catagen, mid-catagen) was determined based on well-validated morphological assessment criteria.

### Chemical stimulation of hair follicles

Chemical reagents were purchased from MedChemExpress. After 24 h of pre-incubation to stabilize follicle function, the culture medium was replaced, and HFs were randomly assigned to treatment groups according to the experimental design: vehicle control (0.1% v/v DMSO), PTN (2.5 ng/ml), Testosterone (10 nM). Culture medium with the corresponding treatments was replenished daily to maintain consistent drug exposure. Follicles were maintained in culture for 2–6 days, after which hair shaft elongation and hair cycle stage were evaluated.

### Data analysis and statistics

#### scRNA-seq raw data processing, integration, clustering and annotation Raw data processing

Raw sequencing paired reads were filtered based on quality and then aligned to the hg38 genome using STAR (v2.7.1a)^75^. To select valid barcode beads, we employed the EmptyDrops algorithm^76^, calculating the cosine similarity between beads based on their oligo types and counts to analyze the raw beads-by-genes matrix. After merging the valid beads within the same cell, a cell versus gene UMI count matrix was generated using PISA (v0.2)^77^.

### Ambient RNA removal

To remove ambient RNA contamination from our single-cell RNA-seq data, we employed SoupX (v1.6.2)^78^. First, we created a SoupChannel object using the raw and filtered matrices. We then set the clusters in the SoupChannel object and estimated the contamination using the autoEstCont function. Finally, we adjusted the counts to remove ambient RNA contamination using the adjustCounts function, rounding the results to integers.

### Doublet removal and quality control

To detect doublets, we used the scDblFinder (v1.12.0)^79^ software. We first created a SingleCellExperiment object using raw count data for each library. We then ran the scDblFinder function with the parameters: clusters=TRUE, aggregateFeatures=TRUE, nfeatures=25, and processing="normFeatures" for each library. The doublet detection results were added to the Seurat object for further analysis. To ensure data quality, we performed an initial filtering step to remove cells with a low UMI count (UMI < 100) for each sample. We then merged all samples into one Seurat object and calculated the median and Median Absolute Deviation (MAD) of these metrics for all cells. Values deviating more than four MADs from the median were flagged as outliers. Finally, we removed cells identified as outliers and doublets from all sample libraries.

### Data integration, clustering and annotation

To merge scRNA-seq data from multiple libraries and mitigate batch effects, we performed an integrated analysis using Seurat (v4.3.0)^80^ in R. This involved identifying variable features across libraries, detecting anchors between datasets, integrating data using Robust Principal Component Analysis (RPCA) and performing dimensionality reduction. Cells were clustered using a resolution of 0.8, and the first 30 principal components (PCs) were used to generate a Uniform Manifold Approximation and Projection (UMAP) plot for data visualization. Cell clusters were assigned cell type annotations based on the expression of established marker genes **(Extended Data Fig. 2b)**.

In order to further analyze the hDSC and HFSC lineages, we extracted the corresponding cell types from each lineage and re-integrated the cells for subsequent pseudotime analysis. The integration of the HFSC lineage was re-integrated using Canonical Correlation Analysis (CCA), and the hDSC lineage was re-integrated using RPCA.

To identify the subpopulations of the HFSC, we specifically selected HO samples corresponding to the age groups of AGA patients. aHFSC and qHFSC were isolated from these samples. Next, we employed CCA to re-integrate the HFSC and subsequently re-clustered these cells. By examining differential gene expression and cell clusters, dqHFSC was defined based on the original qHFSC population.

### Integration with mouse hfDSC data

The GSE115424 dataset was downloaded from the GEO database (https://www.ncbi.nlm.nih.gov/geo/)^13^. In line with the cell quality control standards, the filtered mouse dataset was integrated with human dataset by using the RPCA method in Seurat (v4.3.0). Cell type annotations were made based on reported marker genes. Subsequently, the DP, DS, and hfDSC/hDSC were extracted to run UMAP for data visualization.

### scATAC-seq raw data processing, integration, clustering and annotation Raw data processing

Briefly, raw sequencing reads were aligned to the hg38 genome using Chromap (v0.2.1)^81^. Barcode beads with fragment counts below the knee point were considered empty droplets and removed to ensure valid cell capture. To address multiple beads within a single droplet, we used d2c (v1.5.3)^82^ to merge barcode beads based on fragment distribution similarity. Following bead merging, MACS2 (v2.28)^83^ was employed to identify enriched genomic regions (peaks), resulting in a cell-by-peak matrix for each library.

### Doublet removal and quality control

We employed the scDblFinder (v1.12.0)^79^ software to detect doublets in our scATAC-seq data. First, we generated a union peak set by merging peaks from multiple scATAC-seq sample libraries and filtering out those with inappropriate lengths. This resulted in a cell-by-union peak matrix for the scATAC-seq dataset. We then analyzed the scATAC-seq matrix for each library using the scDblFinder (v1.12.0)^79^ function with the following parameters: clusters=TRUE, aggregateFeatures=TRUE, nfeatures=25, and processing=normFeatures.

Additionally, to ensure high-quality scATAC-seq data, we removed cells that did not meet the following thresholds: nCount_ATAC > 2000, nCount_ATAC < 20000, FRiP > 0.3, blacklist_fraction < 0.1, nucleosome_signal < 1, and TSS.enrichment > 2.

### Data integration, clustering and cell type annotation

For processing scATAC-seq data, we applied TF-IDF normalization and employed Singular Value Decomposition (SVD) for data dimensionality reduction by R package Signac (v1.6.0)^84^. To mitigate batch effects, we utilized the Signac (v1.6.0) for integrating scATAC-seq data. The cell clustering method used was consistent with that of scRNA-seq. Subsequently, we utilized the GeneActivity function in Signac (v1.6.0) to calculate the gene activity scores based on open chromatin regions (peaks). Using the gene activity scores, we integrated scRNA-seq data with scATAC-seq data to identify the scATAC cell types by scRNA cell types projection.

### Stereo-seq raw data processing, cell segmentation, clustering and annotation Raw data processing

We utilized the MGI DNBSEQ-Tx sequencer to generate single-end Stereo-seq fastq files, which contained CID (coordinate identity), MID (molecular identity, UMI), and cDNA sequences. The processing workflow for Stereo-seq raw data was performed using the STOmics Analysis Workflow (SAW v6.0, https://github.com/STOmics/SAW). The obtained reads were aligned to the human reference genome (hg38). Subsequently, low-quality alignments (MAPQ <10) were filtered out. The remaining sequences were annotated with overlapping genes. Finally, the expression matrix containing CID was constructed using the exon reads. The lasso tool in STOmics Cloud (https://cloud.stomics.tech) was used to exclude DNBs without tissue/cells based on the MID density and the staining image.

### Cell segmentation based on image

We followed the method of Wei *et al*., 2022^31^ by overlaying the ssDNA staining image from the same section onto the Stereo-seq chip image to segment cells. First, the Stereo-seq data is converted into a grayscale image, where each pixel represents a DNB. The ssDNA image is then manually registered with the obtained grayscale image to align the DNBs of the staining image with the pixels of the grayscale image. The cell segmentation and tissue region clustering are implemented in the scikit-image package (v0.19.3)^85^. The background noise of the registered ssDNA staining image is filtered using a global threshold. The cell positions are obtained by calculating a Gaussian-weighted local threshold, with a block size of 9 and an offset of 0.03. The cells are segmented using exact Euclidean distance transformation with a minimal distance of 7, and the resulting cell labels are expanded by 2 pixels outward (function expand_labels). The UMI of all DNBs in the corresponding segmentation are merged by genes to obtain a gene-cell expression matrix for downstream analysis.

### Spatially-constrained clustering of Stereo-seq data

After cell segmentation, cells in the raw matrix with UMI > 50 were retained and further normalized using Seurat (v4.3.0.1) SCTransform function. To incorporate spatial information, we utilized Squidpy to combine the spatial k-nearest neighbor graph 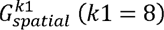 and the k-nearest neighbor graph 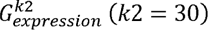 of transcriptome data. The combined graph 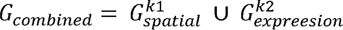 was then subjected to Leiden clustering with a resolution of 5, taking into account the follicle structure to accurately delineate the spatial organization of tissue regions. Hair follicle regions were identified based on anatomical structure^37^.

### Cell type annotation

We employed the robust cell type decomposition (RCTD)^36^ to predict the cell type of Stereo-seq based on scRNA-seq annotations in R spacexr package (v2.2.1). For each cell type in the scRNA-seq data, we down-sampled to 1,000 cells and created a reference object using the Reference function. The query object was constructed using the expression matrix and spatial coordinates of the Stereo-seq data with the SpatialRNA function. Subsequently, we used the reference object to predict the cell type weights for each cell (function create. RCTD, run.RCTD, doublet_mode = doublet). Only cells with successfully predicted cell labels were retained.

### Spatial cell colocalization analysis

Spatial cell colocalization was performed using a K-nearest neighbor (KNN)-based approach with hypergeometric testing. Briefly, for each cell, the *k* nearest spatial neighbors were identified using Squidpy (‘sq.gr.spatial_neighbors’, ‘n_neighs = 10’, ‘coord_type = "generic"’). The resulting spatial adjacency matrix was multiplied by a cell type indicator matrix to count the number of neighboring cells for each cell type pair. For each cell type pair (From→To), enrichment was quantified by log_2_ fold change *log_2_FC = log_2_((x/N)/(n/M))*, where *M* is the total number of neighbor relationships, *N* is the total neighbor count of the source cell type, *n* is the total neighbor count of the target cell type, and *x* is the observed co-occurrence count. Statistical significance was assessed by one-sided hypergeometric test (*P* = 1 − hypergeom.cdf(*x* − 1); *M*, *n*, *N* as defined above). The percentage of target cell type among all neighbors of the source cell type was calculated as *x*/*N*.

Colocalization results were visualized as dot plots. Dot size was scaled by log_10_-transformed co-localized cell counts *Log_10_(Counts)* with larger dots indicating stronger spatial co-occurrence. Only cell type pairs with *P* < 0.05 were displayed.

### RNA velocity analysis

RNA velocity analysis of hDSC and HFSC lineages was performed using Python Dynamo package (v1.3.2)^38^. The raw matrix of unspliced and spliced RNA were obtained from the processed files using SAW (v6.0). After matching cell segmentation labels, gene splicing and degradation rates were estimated to obtain the relative abundance of nascent (unspliced) and mature (spliced) RNA in each cell. Highly expressed genes with significant Moran’s I index were selected as feature genes, and the preprocess_adata function was used to process the original spliced and unspliced matrices. Subsequently, kinetic parameters and gene-wise RNA velocity were estimated and mapped to the spatial positions of the follicles for visualization. A continuous vector field was established using vf function, and spatial pseudotime was obtained through the ddhodge function in the learned vector field, thereby elucidating the process of cell fate changes in the three lineages.

### Monocle 3 analysis

In hair follicle epithelial and mesenchymal cell differentiation lineages, we performed pseudotime analysis based on Monocle 3 (v1.3.1)^86^ using the scRNA-seq raw count matrix extracted from each lineage. Firstly, the count data were standardized and dimensionally reduced using the preprocess_cds function and reduce_dimension function, respectively. Trajectory learning was performed on the re-integrated UMAP of each lineage using the learn_graph function. After selecting hDSC and qHFSC as stem cells respectively, the pseudotime function was used to evaluate the pseudotime of each cell to further validate the conclusions from the RNA velocity analysis.

For the subpopulation transition lineage analysis of HFSC, qHFSC was selected as the starting point of the lineage, and the same analysis methods mentioned above were used to obtain the trajectory of the HFSC subpopulation.

### Monocle 2 analysis

The differentiation trajectories of the human hair follicle epithelial lineages and mouse hair follicle mesenchymal lineages were analyzed using Monocle 2 (v2.22.0)^39^. The top marker genes calculated by the Seurat FindAllMarkers function were input into Monocle 2 to identify the cell differentiation trajectories. Then the standardized data was performed dimensionality reduction via the DDRTree method and visualized with the plot_cell_trajectory function. To identify genes influencing the lineage of the HFSC subpopulation, we used the BEAM function to identify lineage branch-enriched genes and visualized gene expression using R ComplexHeatmap (v2.14.0)^87^.

### Gene regulatory network (GRN) inference using SCENIC+

To delineate the lineage-specific transcriptional regulatory networks driving the divergence of human hair follicle epithelial and mesenchymal compartments, we employed the SCENIC+ pipeline (v1.0a2)^41^. First, scATAC-seq data were processed using pycisTopic (v2.0a0) to identify high-quality accessible regions. Consensus peaks were called using MACS2 (v2.28) with the parameters --nomodel --shift 73 --extsize 146. Subsequently, pycistarget was utilized to perform motif enrichment analysis on these candidate cis-regulatory elements against the v10_nr_macfas_hg38 motif database. Following the establishment of region-to-gene links, we integrated the scRNA-seq expression matrix and the scATAC-seq region accessibility matrix to construct enhancer-driven gene regulatory networks (eRegulons). The SCENIC+ algorithm identified confident TF-Region-Target Gene triplets by assessing the Spearman correlation across TF expression, region accessibility, and target gene expression. Finally, to isolate the key regulators dictating epithelial versus mesenchymal identities, we scored the activity of each eRegulon followed by differential eRegulon activity analysis to prioritize lineage-specific TFs.

### DEGs and GO term enrichment

To identify genes associated with aging and AGA progression, we compared the differentially expressed genes (DEGs) between young adults and elderly adults, as well as between HO and AO, and AF samples. Pseudobulks were generated by calculating the average gene counts for each sample. Subsequently, the samples were grouped according to age or disease status, and the DESeq function (fitType = mean, minReplicatesForReplace = 7) from R DESeq2 (v1.32.0) was used to calculate the differential genes. The DEGs (|log_2_fold change| > 0.5 and *P* < 0.05) obtained from each comparison were input into the Metascape online tool for gene ontology (GO) enrichment analysis (https://metascape.org/gp/index)^88^. The visualization of GO terms was implemented using ComplexHeatmap (v2.14.0)^87^.

### Gene module analysis

To evaluate the spatial expression patterns of distinct cell types in the Stereo-seq data, the top 10 marker genes ranked by log_2_fold change (*P* < 0.05) were selected as marker gene modules and used as input for the Seurat (v4.3.0) AddModuleScore function to assess the spatial characteristics of cell type expression patterns. To investigate the proliferation status of hDSC and qHFSC, we followed the Seurat (v 4.3.0) workflow and utilized the CellCycleScoring function to assess the cell cycle score. Additionally, we examined the aging status of hDSC by evaluating the expression of published aging-related gene modules^53,54^ using the AddModuleScore function. To further validate that the dqHFSC phenotype emerges independently of the hair cycle stage, we calculated module scores using the top 50 dqHFSC signature genes to assess strictly anagen-phase follicles across the HO, AO, and AF samples.

### Cell–cell interaction analysis

CellChat (v.1.6.1)^55^ detects ligand-receptor interactions on the integrated scRNA-seq data according to standard procedures. The single cell expression matrix and cell type information of young and old subjects were imported into CellChat, and the two groups each included 40 cell types. The overall communication probability between cell groups is calculated by using the computeCommunProb function with trim set to 0.1. The filterCommunication function is used to remove communication events that only occur in a few cells and ensure that at least 10 cells are involved in each communication event to improve the reliability of analysis.

### Statistical analysis

R (v 4.2.3) were used for statistical analysis. The statistical results of the data were described in the corresponding figures and legends. For Pearson correlation, the statistical significance of positive correlation or negative correlation is set to *P* < 0.05, the correlation coefficient is greater than 0.4, which is positive correlation, and less than -0.4, which is negative correlation. Shading indicates the 95% confidence interval along the correlation line. In the box plots, the central line represents the median, the box limits represent the upper and lower quartiles, and the points represent the distribution of the samples.

### In-situ hDSC count and dispersion statistic

Cell counting in situ were performed by ImageJ^89^. The counting of hDSC was performed by quantifying SOX2 positive signals. Click on "multi-point" to count all positive cells. The dispersion analysis was performed by ImageJ (National Institutes of Health). The distance between hDSC was analyzed with SOX2 positive pictures. Convert the image to grayscale (8-bit) image, click Analyze, Analyze Particles, get the result XM, YM as the center of mass coordinates. Calculate the distance between two points formula in Excel: point 1 (X1, Y1), point 2 (X2, Y2): SQRT((X1-X2) ^2+(Y1-Y2) ^2). Open Rstudio, import the distance between two points txt. data and calculate the distance between the points in the file.

### Quantitative analysis of immunofluorescence staining-positive cells

Quantitative analysis of target protein-positive cells was performed using ImageJ software (National Institutes of Health, Bethesda, MD, USA). In brief, five randomly selected, non-overlapping high-power fields (HPF, 200× or 400× magnification) were captured per biological replicate under a fluorescence microscope, with consistent exposure parameters across all experimental groups to avoid imaging bias. For each image, the DAPI-stained total cell number was counted automatically using the ImageJ ‘Analyze Particles’ function after uniform threshold adjustment. The number of target protein-positive cells was quantified manually or by semi-automated thresholding in the corresponding fluorescence channel, with positive cells defined as those displaying fluorescence intensity exceeding the background threshold of negative control samples. The percentage of positive cells was calculated as the ratio of positive-stained cells to total DAPI-positive nucleated cells, and the mean value of each group was determined from at least three independent biological replicates.

## Data availability

All data generated in this study are freely accessible in CNGB Nucleotide Sequence Archive under accession code CNP0006019. All other data are in the main paper or the supplementary materials. Processed data can be interactively explored and downloaded from https://db.cngb.org/stomics/hhaamstar/.

## Code availability

All code used to analyze the data is available online at https://github.com/BGI-DEV-REG/HHAAMSTAR.

## Acknowledgments

Pengcheng Guo was supported by the National Key R&D Program of China (2025YFC3508602), China Postdoctoral Science Foundation (2025M772609) and Natural Science Foundation of Zhejiang Province, China (LMS26H110001); Xiaoyu Wei was supported by the National Natural Science Foundation of China (32300707); Yanwen Xu was supported by the Construction Fund of Key Medical Disciplines of Hangzhou (2025HZZD08) and School of Medicine, Westlake University, Youth Physician-Scientist Cultivation Program (2025SOM01); Mingxing Lei was supported by the National Key Research and Development Program of China (2023YFC2508200), National Natural Science Foundation of China (82373509, 82574005); Zhenxing Wang was supported by the National Natural Science Foundation of China (82322046); Pengfei Cai was supported by the National Natural Science Foundation of China (32400550) and Tao Yang was supported by Guangdong Genomics Data Center (2021B1212100001). We sincerely thank the China National GeneBank (CNGB) and Cocalero Biotech Co., Ltd. for the support they provided. We would like to thank DCS Cloud (https://cloud.stomics.tech/) for providing the computational resources and software support necessary for this study.

## Author contributions

X.W., J.Z., P.G., M.L., L.L., X.X., Z.W., and D.N. conceived the idea; J.Z., P.G., M.L., L.L., X.X., Z.W., and D.N. supervised the work; X.W., Y.X., P.G., Z.Z., Y.Yu., and H.C. designed the experiments; Y.X., R.L., Y.Yu., Z.Y. and X.L. performed the majority of the experiments with the help from Y.J., F.Z., H.S., Y.Yuan, Y.H., and J.D., R.L. performed data analysis with the help from P.C and S.W.; T.Y. performed database construction; Y.G., H.L., Y.L., J.X. and S.W. gave relevant advice; X.W., P.G. and Y.X wrote the manuscript with input from all authors.

## Correspondence and requests for materials

should be addressed to Pengcheng Guo.

## Supplementary table legends

**Supplementary Tables 1–16**

**Supplementary Table 1 | scRNA-seq data quality control.**

**Supplementary Table 2 | scATAC-seq data quality control.**

**Supplementary Table 3 | Stereo-seq data quality control.**

**Supplementary Table 4 | Top 20 marker genes for 40 cell types identified by scRNA-seq.**

**Supplementary Table 5 | Transcriptional regulatory networks of hair follicle epithelial lineages.**

**Supplementary Table 6 | Transcriptional regulatory networks of hair follicle mesenchymal lineages.**

**Supplementary Table 7 | DEGs between young and old in hDSC.**

**Supplementary Table 8 | GO terms of hDSC DEGs between young and old.**

**Supplementary Table 9 | Comparative ligand-receptor interactome of young and aged hair follicle niches.**

**Supplementary Table 10 | DEGs of hDSC in AO vs HO and AF vs HO.**

**Supplementary Table 11 | GO terms of DEGs in hDSC in AO vs HO and AF vs HO.**

**Supplementary Table 12 | DEGs of qHFSC in AO vs HO and AF vs HO.**

**Supplementary Table 13 | GO terms of DEGs in qHFSC in AO vs HO and AF vs HO.**

**Supplementary Table 14 | GO terms of marker genes in aHFSC, dqHFSC and qHFSC.**

**Supplementary Table 15 | Comparative ligand-receptor interactome of HO, AO and AF hair follicle niches.**

## References

1. Clevers, H. (2015). What is an adult stem cell? 350, 1319–1320. doi:10.1126/science.aad7016.

2. Orford, K.W., and Scadden, D.T. (2008). Deconstructing stem cell self-renewal: genetic insights into cell-cycle regulation. Nature Reviews Genetics 9, 115–128. 10.1038/nrg2269.

3. de Morree, A., and Rando, T.A. (2023). Regulation of adult stem cell quiescence and its functions in the maintenance of tissue integrity. Nature Reviews Molecular Cell Biology 24, 334–354. 10.1038/s41580-022-00568-6.

4. Nombela-Arrieta, C., Ritz, J., and Silberstein, L.E. (2011). The elusive nature and function of mesenchymal stem cells. Nature Reviews Molecular Cell Biology 12, 126–131. 10.1038/nrm3049.

5. Hanseul Yang, R.C.A., Yejing Ge, Zhong L. Hua, Elaine Fuchs (2017). Epithelial-Mesenchymal Micro-niches Govern Stem Cell Lineage Choices. Cell. 10.1016/j.cell.2017.03.038.

6. Chen, C.-C., Wang, L., Plikus, Maksim V., Jiang, Ting X., Murray, Philip J., Ramos, R., Guerrero-Juarez, Christian F., Hughes, Michael W., Lee, Oscar K., Shi, S., et al. (2015). Organ-Level Quorum Sensing Directs Regeneration in Hair Stem Cell Populations. Cell 161, 277–290. 10.1016/j.cell.2015.02.016.

7. Chuong, C.-M. (2011). Self-Organizing and Stochastic Behaviors During the Regeneration of Hair Stem Cells. Science.

8. Li Wang, J.A.S., Robin D. Dowell, Rui Yi (2016). Foxc1 reinforces quiescence in self-renewing hair follicle stem cells. Science 351, 613–617. 10.1126/science.aad5440.

9. Joost, S., Annusver, K., Jacob, T., Sun, X., Dalessandri, T., Sivan, U., Sequeira, I., Sandberg, R., and Kasper, M. (2020). The Molecular Anatomy of Mouse Skin during Hair Growth and Rest. Cell Stem Cell 26, 441–457.e447. 10.1016/j.stem.2020.01.012.

10. Jaks, V., Barker, N., Kasper, M., van Es, J.H., Snippert, H.J., Clevers, H., and Toftgård, R. (2008). Lgr5 marks cycling, yet long-lived, hair follicle stem cells. Nature Genetics 40, 1291–1299. 10.1038/ng.239.

11. Zhang, B., and Chen, T. (2023). Local and systemic mechanisms that control the hair follicle stem cell niche. Nature Reviews Molecular Cell Biology 25, 87–100. 10.1038/s41580-023-00662-3.

12. Rahmani, W., Abbasi, S., Hagner, A., Raharjo, E., Kumar, R., Hotta, A., Magness, S., Metzger, D., and Biernaskie, J. (2014). Hair Follicle Dermal Stem Cells Regenerate the Dermal Sheath, Repopulate the Dermal Papilla, and Modulate Hair Type. Developmental Cell 31, 543–558. 10.1016/j.devcel.2014.10.022.

13. Shin, W., Rosin, N.L., Sparks, H., Sinha, S., Rahmani, W., Sharma, N., Workentine, M., Abbasi, S., Labit, E., Stratton, J.A., and Biernaskie, J. (2020). Dysfunction of Hair Follicle Mesenchymal Progenitors Contributes to Age-Associated Hair Loss. Developmental Cell 53, 185–198.e187. 10.1016/j.devcel.2020.03.019.

14. Biernaskie, J., Paris, M., Morozova, O., Fagan, B.M., Marra, M., Pevny, L., and Miller, F.D. (2009). SKPs Derive from Hair Follicle Precursors and Exhibit Properties of Adult Dermal Stem Cells. Cell Stem Cell 5, 610–623. 10.1016/j.stem.2009.10.019.

15. Greco, V., Chen, T., Rendl, M., Schober, M., Pasolli, H.A., Stokes, N., dela Cruz-Racelis, J., and Fuchs, E. (2009). A Two-Step Mechanism for Stem Cell Activation during Hair Regeneration. Cell Stem Cell 4, 155-169. 10.1016/j.stem.2008.12.009.

16. Ramos, R., Swedlund, B., Ganesan, A.K., Morsut, L., Maini, P.K., Monuki, E.S., Lander, A.D., Chuong, C.-M., and Plikus, M.V. (2024). Parsing patterns: Emerging roles of tissue self-organization in health and disease. Cell 187, 3165–3186. 10.1016/j.cell.2024.05.016.

17. Hsu, Y.-C., Li, L., and Fuchs, E. (2014). Transit-Amplifying Cells Orchestrate Stem Cell Activity and Tissue Regeneration. Cell 157, 935–949. 10.1016/j.cell.2014.02.057.

18. Adam, R.C., Yang, H., Ge, Y., Lien, W.-H., Wang, P., Zhao, Y., Polak, L., Levorse, J., Baksh, S.C., Zheng, D., and Fuchs, E. (2018). Temporal Layering of Signaling Effectors Drives Chromatin Remodeling during Hair Follicle Stem Cell Lineage Progression. Cell Stem Cell 22, 398–413.e397. 10.1016/j.stem.2017.12.004.

19. Oh, J.W., Kloepper, J., Langan, E.A., Kim, Y., Yeo, J., Kim, M.J., Hsi, T.-C., Rose, C., Yoon, G.S., Lee, S.-J., et al. (2016). A Guide to Studying Human Hair Follicle Cycling In Vivo. Journal of Investigative Dermatology 136, 34–44. 10.1038/jid.2015.354.

20. Lee, J.H., and Choi, S. (2024). Deciphering the molecular mechanisms of stem cell dynamics in hair follicle regeneration. Experimental & Molecular Medicine 56, 110–117. 10.1038/s12276-023-01151-5.

21. Ji, S., Zhu, Z., Sun, X., and Fu, X. (2021). Functional hair follicle regeneration: an updated review. Signal Transduction and Targeted Therapy 6, 66. 10.1038/s41392-020-00441-y.

22. Lolli, F., Pallotti, F., Rossi, A., Fortuna, M.C., Caro, G., Lenzi, A., Sansone, A., and Lombardo, F. (2017). Androgenetic alopecia: a review. Endocrine 57, 9–17. 10.1007/s12020-017-1280-y.

23. Zhang, C., Wang, D., Wang, J., Wang, L., Qiu, W., Kume, T., Dowell, R., and Yi, R. (2021). Escape of hair follicle stem cells causes stem cell exhaustion during aging. Nature Aging 1, 889–903. 10.1038/s43587-021-00103-w.

24. Zhang, C., Wang, D., Dowell, R., and Yi, R. (2023). Single cell analysis of transcriptome and open chromatin reveals the dynamics of hair follicle stem cell aging. Frontiers in Aging 4. 10.3389/fragi.2023.1192149.

25. Li, G., Yang, L., Duan, S., Chen, M., Zhang, Y., Liu, F., Wang, Y., Li, J., Xu, S., Wu, Z., et al. (2026). Single-cell transcriptomics reveals hair growth retardation mediated by aberrant connective tissue sheath contraction in male androgenetic alopecia. Nat Commun 17. 10.1038/s41467-026-70153-4.

26. Zou, Z., Long, X., Zhao, Q., Zheng, Y., Song, M., Ma, S., Jing, Y., Wang, S., He, Y., Esteban, C.R., et al. (2021). A Single-Cell Transcriptomic Atlas of Human Skin Aging. Developmental Cell 56, 383–397.e388. 10.1016/j.devcel.2020.11.002.

27. Yu, G.T., Ganier, C., Allison, D.B., Tchkonia, T., Khosla, S., Kirkland, J.L., Lynch, M.D., and Wyles, S.P. (2024). Mapping epidermal and dermal cellular senescence in human skin aging. Aging Cell, e14358. 10.1111/acel.14358.

28. Ober-Reynolds, B., Wang, C., Ko, J.M., Rios, E.J., Aasi, S.Z., Davis, M.M., Oro, A.E., and Greenleaf, W.J. (2023). Integrated single-cell chromatin and transcriptomic analyses of human scalp identify gene-regulatory programs and critical cell types for hair and skin diseases. Nature Genetics 55, 1288–1300. 10.1038/s41588-023-01445-4.

29. Wu, S., Yu, Y., Liu, C., Zhang, X., Zhu, P., Peng, Y., Yan, X., Li, Y., Hua, P., Li, Q., et al. (2022). Single-cell transcriptomics reveals lineage trajectory of human scalp hair follicle and informs mechanisms of hair graying. Cell Discovery 8, 49. 10.1038/s41421-022-00394-2.

30. Chen, A., Liao, S., Cheng, M., Ma, K., Wu, L., Lai, Y., Qiu, X., Yang, J., Xu, J., Hao, S., et al. (2022). Spatiotemporal transcriptomic atlas of mouse organogenesis using DNA nanoball-patterned arrays. Cell 185, 1777–1792 e1721. 10.1016/j.cell.2022.04.003.

31. Wei, X., Fu, S., Li, H., Liu, Y., Wang, S., Feng, W., Yang, Y., Liu, X., Zeng, Y.-Y., Cheng, M., et al. (2022). Single-cell Stereo-seq reveals induced progenitor cells involved in axolotl brain regeneration. Science 377, eabp9444. 10.1126/science.abp9444.

32. Yi, R. (2017). Concise Review: Mechanisms of Quiescent Hair Follicle Stem Cell Regulation. Stem Cells 35, 2323–2330. 10.1002/stem.2696.

33. Tierney, M.T., Polak, L., Yang, Y., Abdusselamoglu, M.D., Baek, I., Stewart, K.S., and Fuchs, E. (2024). Vitamin A resolves lineage plasticity to orchestrate stem cell lineage choices. Science 383, eadi7342. 10.1126/science.adi7342.

34. Wang, L., Siegenthaler, J.A., Dowell, R.D., and Yi, R. (2016). Foxc1 reinforces quiescence in self-renewing hair follicle stem cells. Science 351, 613–617. 10.1126/science.aad5440.

35. Niiyama, S., Ishimatsu-Tsuji, Y., Nakazawa, Y., Yoshida, Y., Soma, T., Ideta, R., Mukai, H., and Kishimoto, J. (2018). Gene Expression Profiling of the Intact Dermal Sheath Cup of Human Hair Follicles. Acta Dermato Venereologica 98, 694–698. 10.2340/00015555-2949.

36. Cable, D.M., Murray, E., Zou, L.S., Goeva, A., Macosko, E.Z., Chen, F., and Irizarry, R.A. (2021). Robust decomposition of cell type mixtures in spatial transcriptomics. Nature Biotechnology 40, 517–526. 10.1038/s41587-021-00830-w.

37. Harland, D.P. (2018). Introduction to Hair Development. In The Hair Fibre: Proteins, Structure and Development, pp. 89–96. 10.1007/978-981-10-8195-8_8.

38. Qiu, X., Zhang, Y., Martin-Rufino, J.D., Weng, C., Hosseinzadeh, S., Yang, D., Pogson, A.N., Hein, M.Y., Hoi Min, K., Wang, L., et al. (2022). Mapping transcriptomic vector fields of single cells. Cell 185, 690–711.e645. 10.1016/j.cell.2021.12.045.

39. Qiu, X., Mao, Q., Tang, Y., Wang, L., Chawla, R., Pliner, H.A., and Trapnell, C. (2017). Reversed graph embedding resolves complex single-cell trajectories. Nature Methods 14, 979–982. 10.1038/nmeth.4402.

40. Ma, S., Zhang, B., LaFave, L.M., Earl, A.S., Chiang, Z., Hu, Y., Ding, J., Brack, A., Kartha, V.K., Tay, T., et al. (2020). Chromatin Potential Identified by Shared Single-Cell Profiling of RNA and Chromatin. Cell 183, 1103–1116 e1120. 10.1016/j.cell.2020.09.056.

41. Bravo González-Blas, C., De Winter, S., Hulselmans, G., Hecker, N., Matetovici, I., Christiaens, V., Poovathingal, S., Wouters, J., Aibar, S., and Aerts, S. (2023). SCENIC+: single-cell multiomic inference of enhancers and gene regulatory networks. Nature Methods 20, 1355–1367. 10.1038/s41592-023-01938-4.

42. Bhattacharya, N., Indra, A.K., and Ganguli-Indra, G. (2022). Selective Ablation of BCL11A in Epidermal Keratinocytes Alters Skin Homeostasis and Accelerates Excisional Wound Healing In Vivo. Cells 11. 10.3390/cells11132106.

43. Folgueras, Alicia R., Guo, X., Pasolli, H.A., Stokes, N., Polak, L., Zheng, D., and Fuchs, E. (2013). Architectural Niche Organization by LHX2 Is Linked to Hair Follicle Stem Cell Function. Cell Stem Cell 13, 314–327. 10.1016/j.stem.2013.06.018.

44. Adam, R.C., Yang, H., Ge, Y., Infarinato, N.R., Gur-Cohen, S., Miao, Y., Wang, P., Zhao, Y., Lu, C.P., Kim, J.E., et al. (2020). NFI transcription factors provide chromatin access to maintain stem cell identity while preventing unintended lineage fate choices. Nature Cell Biology 22, 640–650. 10.1038/s41556-020-0513-0.

45. Romano, R.-A., Smalley, K., Liu, S., and Sinha, S. (2010). Abnormal hair follicle development and altered cell fate of follicular keratinocytes in transgenic mice expressing ΔNp63α. Development 137, 1775–1775. 10.1242/dev.053116.

46. Vanhoutteghem, A., Delhomme, B., Hervé, F., Nondier, I., Petit, J.-M., Araki, M., Araki, K., and Djian, P. (2016). The importance of basonuclin 2 in adult mice and its relation to basonuclin 1. Mechanisms of Development 140, 53–73. 10.1016/j.mod.2016.02.002.

47. Flores, A., Schell, J., Krall, A.S., Jelinek, D., Miranda, M., Grigorian, M., Braas, D., White, A.C., Zhou, J.L., Graham, N.A., et al. (2017). Lactate dehydrogenase activity drives hair follicle stem cell activation. Nature Cell Biology 19, 1017–1026. 10.1038/ncb3575.

48. Hwang, J., Mehrani, T., Millar, S.E., and Morasso, M.I. (2008). Dlx3 is a crucial regulator of hair follicle differentiation and cycling. Development 135, 3149–3159. 10.1242/dev.022202.

49. Yu, N., Song, Z., Zhang, K., and Yang, X. (2017). MAD2B acts as a negative regulatory partner of TCF4 on proliferation in human dermal papilla cells. Scientific Reports 7. 10.1038/s41598-017-10350-w.

50. Hiroyuki Matsumura, Y.M., Nguyen Thanh Binh, Hironobu Morinaga, Makoto Fukuda, Mayumi Ito, Sotaro Kurata, Jan Hoeijmakers, Emi K. Nishimura (2016). Hair follicle aging is driven by transepidermal elimination of stem cells via COL17A1 proteolysis. Science 351, aad4395-4391.

51. Ge, Y., Miao, Y., Gur-Cohen, S., Gomez, N., Yang, H., Nikolova, M., Polak, L., Hu, Y., Verma, A., Elemento, O., et al. (2020). The aging skin microenvironment dictates stem cell behavior. Proceedings of the National Academy of Sciences 117, 5339–5350. 10.1073/pnas.1901720117.

52. Pei, D., Shu, X., Gassama-Diagne, A., and Thiery, J.P. (2019). Mesenchymal– epithelial transition in development and reprogramming. Nature Cell Biology 21, 44-53. 10.1038/s41556-018-0195-z.

53. 53. Saul, D., Kosinsky, R.L., Atkinson, E.J., Doolittle, M.L., Zhang, X., LeBrasseur, N.K., Pignolo, R.J., Robbins, P.D., Niedernhofer, L.J., Ikeno, Y., et al. (2022). A new gene set identifies senescent cells and predicts senescence-associated pathways across tissues. Nature Communications 13, 4827. 10.1038/s41467-022-32552-1.

54. Tao, W., Yu, Z., and Han, J.-D.J. (2024). Single-cell senescence identification reveals senescence heterogeneity, trajectory, and modulators. Cell Metabolism 36, 1126–1143.e1125. 10.1016/j.cmet.2024.03.009.

55. Jin, S., Guerrero-Juarez, C.F., Zhang, L., Chang, I., Ramos, R., Kuan, C.-H., Myung, P., Plikus, M.V., and Nie, Q. (2021). Inference and analysis of cell-cell communication using CellChat. Nature Communications 12, 1088. 10.1038/s41467-021-21246-9.

56. Seo, C.H., Kwack, M.H., Kim, M.K., Kim, J.C., and Sung, Y.K. (2017). Activin A induced signalling controls hair follicle neogenesis. Experimental Dermatology 26, 108–115. 10.1111/exd.13234.

57. Wang, N., Zhang, W.-d., Zhong, Z.-y., Zhou, X.-b., Shi, X.-r., and Wang, X. (2023). FGF7 secreted from dermal papillae cell regulates the proliferation and differentiation of hair follicle stem cell1. Journal of Integrative Agriculture. 10.1016/j.jia.2023.10.012.

58. Choi, S., Zhang, B., Ma, S., Gonzalez-Celeiro, M., Stein, D., Jin, X., Kim, S.T., Kang, Y.-L., Besnard, A., Rezza, A., et al. (2021). Corticosterone inhibits GAS6 to govern hair follicle stem-cell quiescence. Nature 592, 428–432. 10.1038/s41586-021-03417-2.

59. Morinaga, H., Mohri, Y., Grachtchouk, M., Asakawa, K., Matsumura, H., Oshima, M., Takayama, N., Kato, T., Nishimori, Y., Sorimachi, Y., et al. (2021). Obesity accelerates hair thinning by stem cell-centric converging mechanisms. Nature 595, 266–271. 10.1038/s41586-021-03624-x.

60. Deng, Y., Wang, M., He, Y., Liu, F., Chen, L., and Xiong, X. (2023). Cellular Senescence: Ageing and Androgenetic Alopecia. Dermatology 239, 533–541. 10.1159/000530681.

61. Krefft-Trzciniecka, K., Piętowska, Z., Nowicka, D., and Szepietowski, J.C. (2023). Human Stem Cell Use in Androgenetic Alopecia: A Systematic Review. Cells 12, 951. 10.3390/cells12060951.

62. Khunkhet, S., Chanprapaph, K., Rutnin, S., and Suchonwanit, P. (2021). Histopathological Evidence of Occipital Involvement in Male Androgenetic Alopecia. Frontiers in Medicine 8. 10.3389/fmed.2021.790597.

63. Takubo, K., Nagamatsu, G., Kobayashi, Chiharu I., Nakamura-Ishizu, A., Kobayashi, H., Ikeda, E., Goda, N., Rahimi, Y., Johnson, Randall S., Soga, T., et al. (2013). Regulation of Glycolysis by Pdk Functions as a Metabolic Checkpoint for Cell Cycle Quiescence in Hematopoietic Stem Cells. Cell Stem Cell 12, 49–61. 10.1016/j.stem.2012.10.011.

64. Tang, Y., Luo, B., Deng, Z., Wang, B., Liu, F., Li, J., Shi, W., Xie, H., Hu, X., and Li, J. (2016). Mitochondrial aerobic respiration is activated during hair follicle stem cell differentiation, and its dysfunction retards hair regeneration. PeerJ 4, e1821. 10.7717/peerj.1821.

65. Sun, P., Wang, Z., Li, S., Yin, J., Gan, Y., Liu, S., Lin, Z., Wang, H., Fan, Z., Qu, Q., et al. (2024). Autophagy induces hair follicle stem cell activation and hair follicle regeneration by regulating glycolysis. Cell & Bioscience 14, 6. 10.1186/s13578-023-01177-2.

66. Choi, N., Kim, W.S., Oh, S.H., and Sung, J.H. (2020). Epiregulin promotes hair growth via EGFR medicated epidermal and ErbB4 mediated dermal stimulation. Cell Proliferation 53, e12881. 10.1111/cpr.12881.

67. Lu, Q., Gao, Y., Fan, Z., Xiao, X., Chen, Y., Si, Y., Kong, D., Wang, S., Liao, M., Chen, X., et al. (2021). Amphiregulin promotes hair regeneration of skin derived precursors via the PI3K and MAPK pathways. Cell Proliferation 54, e13106. 10.1111/cpr.13106.

68. Amberg, N., Sotiropoulou, P.A., Heller, G., Lichtenberger, B.M., Holcmann, M., Camurdanoglu, B., Baykuscheva-Gentscheva, T., Blanpain, C., and Sibilia, M. (2019). EGFR Controls Hair Shaft Differentiation in a p53-Independent Manner. iScience 15, 243–256. 10.1016/j.isci.2019.04.018.

69. Ito, M., Yang, Z., Andl, T., Cui, C., Kim, N., Millar, S.E., and Cotsarelis, G. (2007). Wnt-dependent de novo hair follicle regeneration in adult mouse skin after wounding. Nature 447, 316–320. 10.1038/nature05766.

70. Trueb, R.M. (2002). Molecular mechanisms of androgenetic alopecia. Experimental Gerontology 37, 981–990.

71. Liu, Y., Tosti, A., Wang, E.C.E., Heilmann-Heimbach, S., Aguh, C., Jimenez, F., Lin, S.-J., Kwon, O., and Plikus, M.V. (2025). Androgenetic alopecia. Nature Reviews Disease Primers 11. 10.1038/s41572-025-00656-9.

72. Wang, E.C.E., Dai, Z., Ferrante, A.W., Drake, C.G., and Christiano, A.M. (2019). A Subset of TREM2+ Dermal Macrophages Secretes Oncostatin M to Maintain Hair Follicle Stem Cell Quiescence and Inhibit Hair Growth. Cell Stem Cell 24, 654–669.e656. 10.1016/j.stem.2019.01.011.

73. Lee, J.-H., Tammela, T., Hofree, M., Choi, J., Marjanovic, N.D., Han, S., Canner, D., Wu, K., Paschini, M., Bhang, D.H., et al. (2017). Anatomically and Functionally Distinct Lung Mesenchymal Populations Marked by Lgr5 and Lgr6. Cell 170, 1149–1163.e1112. 10.1016/j.cell.2017.07.028.

74. Yuan, Z., Chen, J., Xu, Y., Zhou, Z., Cai, P., Wei, X., Zheng, H., Zhang, J., Yuan, Y., and Liu, C. (2024). Protocol for optimized dissociation of human scalp tissue for hair follicle transcriptomics by scRNA-seq. STAR Protocols 5, 102848. 10.1016/j.xpro.2024.102848.

75. Dobin, A., Davis, C.A., Schlesinger, F., Drenkow, J., Zaleski, C., Jha, S., Batut, P., Chaisson, M., and Gingeras, T.R. (2013). STAR: ultrafast universal RNA-seq aligner. Bioinformatics 29, 15–21. 10.1093/bioinformatics/bts635.

76. Lun, A.T.L., Riesenfeld, S., Andrews, T., Dao, T.P., Gomes, T., and Marioni, J.C. (2019). EmptyDrops: distinguishing cells from empty droplets in droplet-based single-cell RNA sequencing data. Genome Biology 20, 63. 10.1186/s13059-019-1662-y.

77. Shi, Q., Liu, S., Kristiansen, K., Liu, L., and Mathelier, A. (2022). The FASTQ+ format and PISA. Bioinformatics 38, 4639–4642. 10.1093/bioinformatics/btac562.

78. Young, M.D., and Behjati, S. (2020). SoupX removes ambient RNA contamination from droplet-based single-cell RNA sequencing data. GigaScience 9, giaa151. 10.1093/gigascience/giaa151.

79. Germain, P.-L., Lun, A., Garcia Meixide, C., Macnair, W., and Robinson, M.D. (2022). Doublet identification in single-cell sequencing data using scDblFinder. F1000Research 10, 979. 10.12688/f1000research.73600.2.

80. Hao, Y., Hao, S., Andersen-Nissen, E., Mauck, W.M., 3rd, Zheng, S., Butler, A., Lee, M.J., Wilk, A.J., Darby, C., Zager, M., et al. (2021). Integrated analysis of multimodal single-cell data. Cell 184, 3573-3587 e3529. 10.1016/j.cell.2021.04.048.

81. Zhang, H., Song, L., Wang, X., Cheng, H., Wang, C., Meyer, C.A., Liu, T., Tang, M., Aluru, S., Yue, F., et al. (2021). Fast alignment and preprocessing of chromatin profiles with Chromap. Nature Communications 12, 6566. 10.1038/s41467-021-26865-w.

82. Lareau, C.A., Ma, S., Duarte, F.M., and Buenrostro, J.D. (2020). Inference and effects of barcode multiplets in droplet-based single-cell assays. Nature Communications 11, 866. 10.1038/s41467-020-14667-5.

83. Zhang, Y., Liu, T., Meyer, C.A., Eeckhoute, J., Johnson, D.S., Bernstein, B.E., Nusbaum, C., Myers, R.M., Brown, M., Li, W., and Liu, X.S. (2008). Model-based Analysis of ChIP-Seq (MACS). Genome Biology 9, R137. 10.1186/gb-2008-9-9-r137.

84. Stuart, T., Srivastava, A., Madad, S., Lareau, C.A., and Satija, R. (2021). Single-cell chromatin state analysis with Signac. Nature Methods 18, 1333–1341. 10.1038/s41592-021-01282-5.

85. van der Walt, S., Schönberger, J.L., Nunez-Iglesias, J., Boulogne, F., Warner, J.D., Yager, N., Gouillart, E., and Yu, T. (2014). scikit-image: image processing in Python. PeerJ 2, e453. 10.7717/peerj.453.

86. Cao, J., Spielmann, M., Qiu, X., Huang, X., Ibrahim, D.M., Hill, A.J., Zhang, F., Mundlos, S., Christiansen, L., Steemers, F.J., et al. (2019). The single-cell transcriptional landscape of mammalian organogenesis. Nature 566, 496–502. 10.1038/s41586-019-0969-x.

87. Gu, Z. (2022). Complex heatmap visualization. iMeta 1, e43. 10.1002/imt2.43.

88. Zhou, Y., Zhou, B., Pache, L., Chang, M., Khodabakhshi, A.H., Tanaseichuk, O., Benner, C., and Chanda, S.K. (2019). Metascape provides a biologist-oriented resource for the analysis of systems-level datasets. Nat Commun 10, 1523. 10.1038/s41467-019-09234-6.

89. Collins, T.J. (2018). ImageJ for Microscopy. BioTechniques 43, 25–30. 10.2144/000112517.

